# Simultaneous CRISPR-Cas9-induced double strand breaks are lethal in models of pancreatic cancer

**DOI:** 10.1101/2023.04.03.535384

**Authors:** Selina Shiqing K. Teh, Akhil Kotwal, Alexis Bennett, Eitan Halper-Stromberg, Laura Morsberger, Saum Zamani, Yanan Shi, Alyza Skaist, Qingfeng Zhu, Kirsten Bowland, Hong Liang, Ralph H. Hruban, Chien-Fu Hung, Robert A. Anders, Nicholas J. Roberts, Robert B. Scharpf, Michael Goldstein, Ying S. Zou, James R. Eshleman

## Abstract

While radiation is an effective oncologic therapy, killing cancer by inducing DNA double-strand breaks (DSBs), it lacks specificity for neoplastic cells. We have previously adapted the CRISPR-Cas9 gene-editing technology as a cancer-specific treatment modality targeting somatic mutations in pancreatic cancer (PC). However, its tumoricidal potential remains unclear, especially in comparison to therapeutic doses of radiation. Here, we demonstrate that CRISPR-Cas9-induced DSBs are more cytotoxic in PCs than a comparable number of radiation-induced DSBs. We observed >90% tumor growth inhibition by targeting 9 sites with cancer-specific single-guide RNAs (sgRNAs). Through both bioinformatics and cytogenetics analyses, we found that CRISPR-Cas9-induced DSBs triggered ongoing chromosomal rearrangements, with 87% of structural variants not directly produced from the initial CRISPR-Cas9-induced DSBs, and chromosomal instability (CIN) peaking before cell death. By comparing the cytotoxicity of CRISPR-Cas9- to radiation-induced DSBs, we demonstrate that the number of DSBs required to achieve equitoxic effects was ∼3 times higher for radiation than CRISPR-Cas9. Finally, we show that PC cells that had survived CRISPR-Cas9 targeting retained susceptibility to subsequent CRISPR-Cas9-induced DSBs at different genomic sites with >87% growth inhibition. Together, our data support the therapeutic potential of CRISPR-Cas9 as an anti-cancer strategy.

## Introduction

Chromosomal instability (CIN), a cancer hallmark, produces numerical and/or structural chromosomal abnormalities. Under selective pressure, such as tumor evolution and therapeutic intervention, CIN enables clonal evolution of heterogeneous karyotypes and phenotypic adaptation, promoting drug-resistant clones, bypassing oncogene addiction, and expanding oncogene-independent subclones (1–3). However, CIN carries a fitness cost: many karyotypes are non-viable, and extreme CIN, intolerable to cancer cells, correlates with better prognosis (4, 5). Elevated chromosome mis-segregation that drives high CIN can suppress tumor growth (6–8), suggesting that for tumors to propagate sustainably, the equilibrium between the tumor-promoting CIN and tumor-suppressive CIN has to be optimal, and that strategies to shift the equilibrium towards cancer cell killing could be leveraged to improve patient outcomes (1).

Double-strand DNA breaks (DSBs) are known to be the most lethal of all DNA lesions (9, 10). This cytotoxicity originates from the DNA damage response that either repairs DSBs or triggers senescence or cell death, and many conventional anti-cancer strategies, such as radiation therapy and some chemotherapies, exploit it for cancer killing (9). Despite multiple repair pathways, DSBs remain dangerous to cell survival as they directly disrupt the genome integrity, and erroneous repair can fuel CIN (9–11). One such consequence is the initiation of breakage-fusion-bridge (BFB) cycles, in which broken chromosomes fuse to form dicentrics, anaphase breaks, SVs, and telomere-free chromosome ends, propagating further genomic rearrangements (12–14).

Studies using endonucleases, such as I-*Sce*I and CRISPR-Cas9 that can induce DSBs at specific locations, have shown a positive correlation between increased number of DSBs and growth inhibition (15–17). CRISPR-Cas9-induced DSBs have been linked to chromothripsis (15), chromosomal rearrangements (18, 19), and cytotoxicity (20–22). However, most studies seek to minimize cytotoxicity to improve editing (22–24), with very few exploiting this cytotoxicity for cell killing (20, 25, 26). Recent studies have also reported upregulation of p53 pathway in Cas9-expressing cells to inhibit genetic perturbations, indicating *TP53* status must be considered when assessing CRISPR-Cas9-induced cytotoxicity (27–29).

We present evidence that CRISPR-Cas9 is a potent anti-cancer strategy for pancreatic cancer (PC). PC is the third leading cause of cancer death in the United States with a dismal 5-year survival rate of 13% (30). More than 90% of PCs are pancreatic ductal adenocarcinomas (PDACs) (31), and approximately 70% of PDACs harbor *TP53* inactivation (32). While radiation therapy damages DNA directly or indirectly via free radicals (33), we find CRISPR-Cas9-induced DSBs are more cytotoxic than clinically relevant doses of radiation, driving extreme CIN and cell death. More broadly, CRISPR-Cas9 may serve as a selective, highly cytotoxic approach against cancers.

## Results

### CRISPR-Cas9-induced DSBs inhibit cancer cell growth

We designed multi-target sgRNAs with 2-16 target sites in non-coding regions of the human genome to avoid confounding cytotoxicity caused by gene essentiality (Supplemental Table 1, sgRNA design previously described (26)). Negative controls included two non-targeting sgRNAs (NT and NT2), while positive controls included three sgRNAs targeting repetitive elements. An *HPRT1*-targeting sgRNA was also designed to validate Cas9 activity via 6-thioguanine resistance. The number of predicted target sites of each sgRNA is noted in parentheses (e.g., 52F(3) has 3 sites; (rep) denotes targets in repetitive elements). We generated Cas9-expressing cell lines from two *TP53*-inactivated PC lines, Panc10.05 (homozygous I255N) and TS0111 (homozygous C275Y) and documented Cas9 activity (Supplemental Figure 1A) using a previously described assay (26). Transduction of negative and positive control sgRNAs into parental, dead Cas9 (dCas9)-expressing, and Cas9-expressing cells showed growth inhibition only in Cas9-expressing cells with positive control, repetitive-element-targeting sgRNAs, indicating multiple DSBs are required for growth inhibition (Supplemental Figure 1B).

To test the hypothesis that growth inhibition increased with the number of simultaneously induced DSBs, we transduced multi-target sgRNAs into Cas9-expressing cells and performed clonogenicity and cell viability assays, ending when non-targeting controls reached full confluence (1-2 months). We found that in general, clonogenic and cell survival decreased as a function of the number of sgRNA target sites (Figure 1, A and B). Multi-target sgRNAs with 12 target sites or more consistently produced >90% clonogenic inhibition, with 230F(12) demonstrating cytotoxicity similar to that of positive control sgRNAs. However, some variability was observed; for example, 451F(6) and 176R(7) sgRNAs were less inhibitory than 715F(5) sgRNA. Significant decrease in clonogenic inhibition was only detected in one of the cell lines treated with the 531F(2) sgRNA (Dunn-Sidak test, *P = 0.040*).

**Figure 1.**
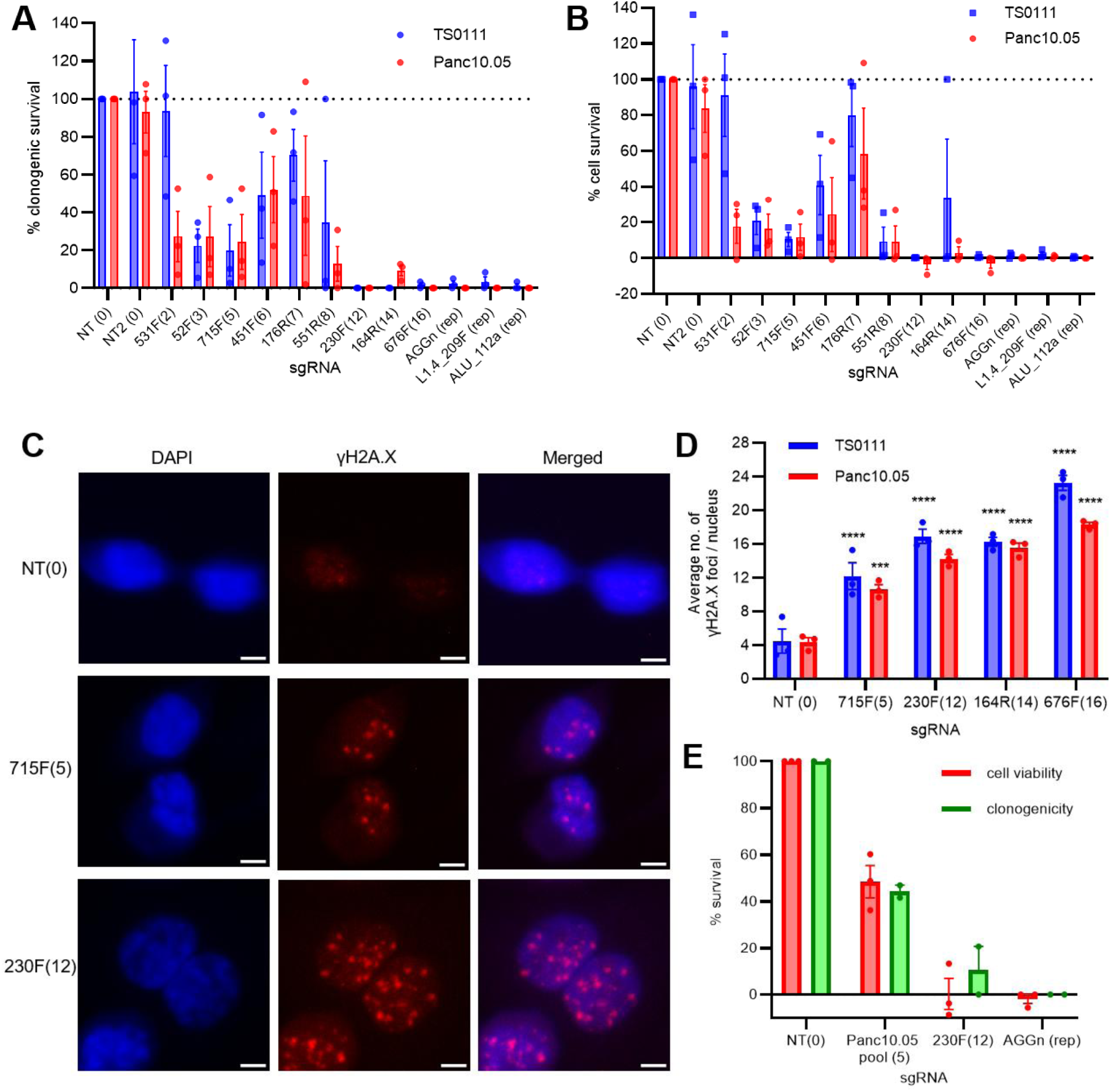
Increased CRISPR-Cas9-induced double strand breaks (DSBs) inhibited cancer cell growth. **(A)** Clonogenic survival with increased number of CRISPR-Cas9 target sites in the human genome of two pancreatic cancer (PC) cell lines. Number of target sites in parentheses, “rep” indicates repetitive element-targeting. N=3; mean ± SEM, normalized to NT. **(B)** Cell survival with increased number of CRISPR-Cas9 target sites as detected by alamarBlue cell viability assay. N=3; mean ± SEM, normalized to NT. **(C)** Representative images of γH2A.X staining in Panc10.05 cells transduced with either non-targeting (NT), 715F(5), or 230F(12) multi-target sgRNAs. Images at 40X magnification; Scale bar is 5µM. N=3. **(D)** Number of γH2A.X foci as a function of the number of CRISPR-Cas9 target sites. >100 nuclei were analyzed for each condition. Dunnett’s test between NT and each multi-target sgRNA; *** *P<0.001,* **** *P<0.0001.* N=3; mean ± SEM. **(E)** Clonogenic and cell survival 21 days after electroporating in CRISPR-Cas9 with multi-target sgRNAs or a pool of 5 sgRNAs targeting different noncoding mutations in the Panc10.05 genome. N=2/3; mean ± SEM, normalized to NT.

We performed γH2A.X staining to quantify CRISPR-Cas9-induced DSBs following multi-target sgRNA transduction. Given the hypodiploidy of Panc10.05 (39-40 chromosomes) and the hyperdiploidy of TS0111 (50–61), we expected more foci in TS0111 at the same target sites. We found that an increased number of sgRNA target sites correlated with an increased number of γH2A.X foci in both cell lines 48 hours post transduction, with TS0111 cells generally exhibited more foci (Figure 1, C and D; Supplemental Figure 1C). Transduction of ALU_112a resulted in an excessive number of γH2A.X foci that was uncountable (Supplemental Figure 1D), consistent with the fact that ALU_112a is expected to generate ∼66,533 DSBs per cell (Cas-OFFinder (34)). γH2A.X staining of AGGn (repetitive element-targeting sgRNA) at earlier time points also showed that foci were below saturation before 48 hours (Supplemental Figure 1E). The baseline foci number was ∼4 (Figure 1D), consistent with previous literature which established that malignant cells tend to have elevated levels of γH2A.X at baseline (35, 36). Since the number of γH2A.X foci correlated with the number of CRISPR-Cas9 target sites and none of the cells transduced with multi-target sgRNAs showed anomalous foci formation, our data suggest that the clonogenic inhibition by multi-target sgRNAs was a result of targeted DSB induction and not widespread off-target activities.

To ensure that our observations were not limited to constitutive expression of CRISPR-Cas9, we electroporated CRISPR-Cas9 ribonucleoprotein (RNP) complex into both cell lines for transient expression. We introduced NT, 230F(12), and AGGn sgRNAs, and for Panc10.05, a combination of 5 sgRNAs targeting 5 different Panc10.05-specific noncoding mutations (Panc10.05 pool, designed using a protospacer adjacent motif (PAM)-based, cancer-specific approach previously described (26); Supplemental Table 2), into cells and performed clonogenic survival assay for 21 days, instead of 1-2 months in our previous assays. Our results were similar to our data in Figure 1 A and B, in which 230F(12) showed >90% clonogenic inhibition (Figure 1E; Supplemental Figure 2A). Differences could be attributed to the confounding effect of cells that did not receive the CRISPR-Cas9 RNPs and survived. Interestingly, the Panc10.05-specific sgRNAs (each at 1/5 of the concentration used for multi-target sgRNAs) yielded >50% growth inhibition (Figure 1E), showing that transient expression of these mutation-targeting sgRNAs can inhibit PC cell growth.

The cell line-specific sgRNAs target somatic, non-coding mutations in cancer cells that are absent in patient-matched normal cells (26). To study the impact of these cancer-specific sgRNAs on normal cells, we treated two primary skin fibroblasts and two cancer-associated fibroblasts cell lines, all derived from patients, with a pool of 9 sgRNAs specific to either TS0111 or Panc10.05, cultured 3-4 weeks, and assessed cell survival (Supplemental Figure 2B; Supplemental Table 2 and 3). No significant differences versus negative controls were observed, indicating these cancer-specific sgRNAs do not inhibit growth of normal cells lacking the targets.

### Simultaneous CRISPR-Cas9 targeting inhibits tumor and metastatic growth

To investigate whether increasing CRISPR-Cas9-induced DSBs would reduce tumor growth, we first transduced Panc10.05 Cas9-expressing cells with NT, 715F(5), 230F(12), ALU_112a, or the 9-sgRNA Panc10.05 pool (as in Supplemental Figure 2B; Supplemental Table 2). After puromycin selection of transduced cells, we injected them subcutaneously into both flanks of nude mice and monitored them for 6 weeks. The percentage of NT tumors present post xenograft peaked at 90%, followed by 30% of 715F(5) and Panc10.05 pool tumors, and 20% of 230F(12) tumors (Figure 2A); ALU_112a yielded no tumors (data not shown). Significant decreases in tumor volumes between NT and 230F(12) or Panc10.05 pool were observed as early as week 4 (Figure 2B; Dunn-Sidak test, NT and 230F(12): *P=0.010*, NT and Panc10.05 pool: *P=0.043*, all N=10). By week 6, significant decreases in tumor volumes between NT and the remaining treatment groups were observed (Figure 2B, all *P<0.0001*). Tumors were harvested and weighed by the end of week 6 (Supplemental Figure 2C), which showed significant decreases in tumor weight between NT and the rest of the treatment groups (all *P<0.0001*), but no differences in body weight (Supplemental Figure 2D). To determine whether CRISPR-Cas9 activity had occurred in tumors harvested, we extracted genomic DNA from the two outlier 230F(12) tumors, PCR amplified the 230F(12) target regions, and performed next generation sequencing (NGS). We found that the mutation frequency was 0.97% and 0.64% in 230F(12) tumors, suggesting insufficient CRISPR-Cas9 activity enabled tumor growth.

**Figure 2.**
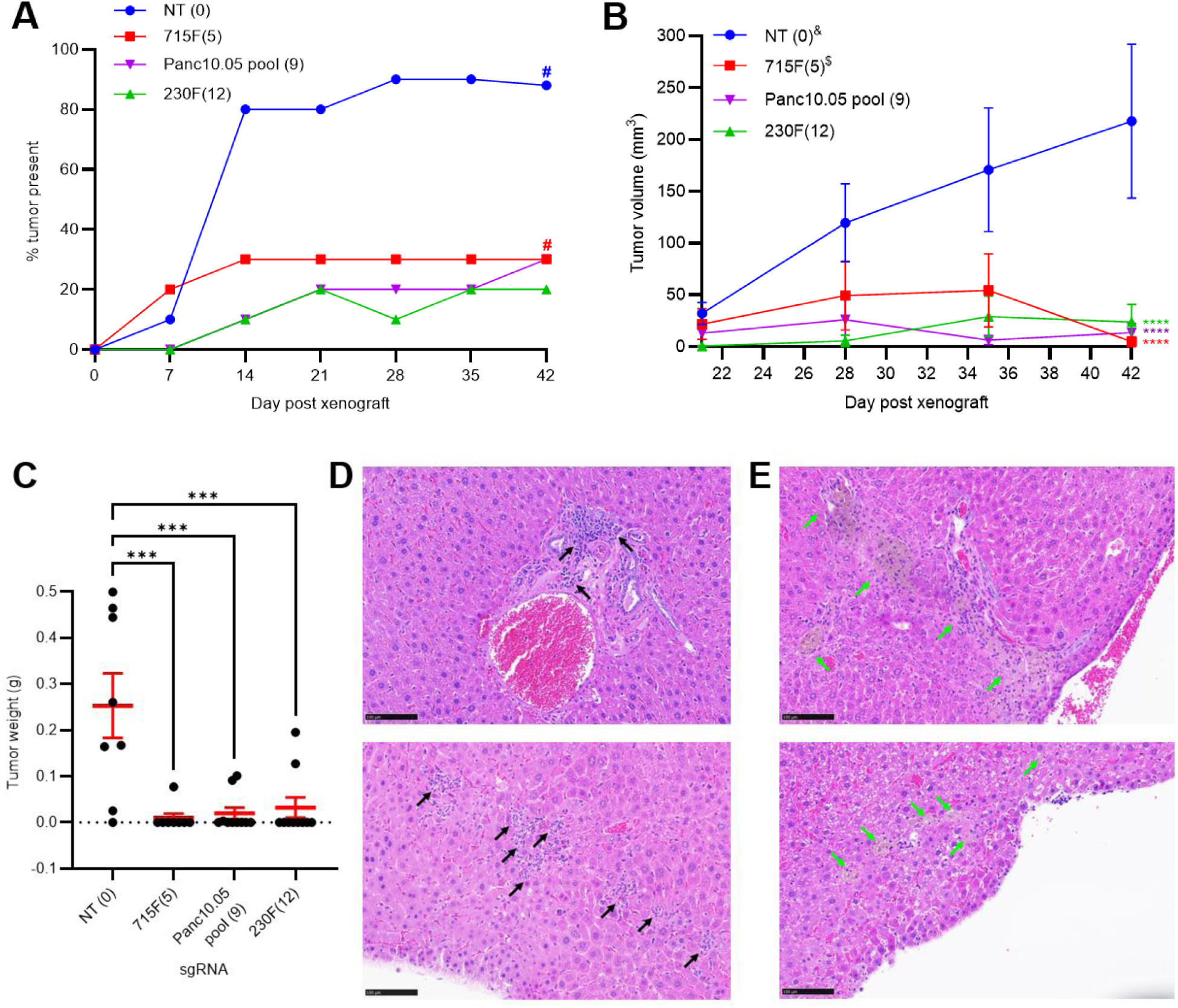
Simultaneous CRISPR-Cas9 targeting inhibited tumor growth. **(A-C)** Tumor growth experiment in subcutaneous xenograft models. Panc10.05 Cas9-expressing cells transduced with the following sgRNAs: NT, 715F(5), 230F(12), or a pool of 9 sgRNAs targeting different noncoding mutations unique to Panc10.05 (Panc10.05 pool) were injected into nude mice for tumor growth. **(A)** Percentage of tumors present post-xenograft. # indicates absence of two data points due to early death around week 5 (33-36 days). **(B)** Tumor volume measurements post-xenograft. Dunn-Sidak test between NT and the other treatment groups on week 6, all *P<0.0001*. N=10; mean ± SEM. & indicates absence of week 5 and 6 data points of two tumors due to early death. $ indicates absence of two data points from week 6 due to early death. **(C)** Tumor weight measurements on week 6 post-xenograft. Dunnett’s test between NT (N=8) and 715F(5): *P=0.0003* (N=8), 230F(12): *P=0.0008* (N=10), and Panc10.05 pool: *P=0.0004* (N=10). Mean ± SEM was shown. **(D-E)** Metastatic growth experiment in hemi-spleen injection mouse models of liver metastasis. Hematoxylin and eosin (H&E) staining of the liver sections of mice treated with **(D)** NT (N=7) or **(E)** 230F(12) (N=5) sgRNA-expressing PC cells. Black arrow: tumor growth; green arrow: tumor regression. The top and bottom panels represent liver sections from two different mice of the same treatment group. Images at 20X magnification; scale bar is 100µM.

Since the primary site of PC metastasis is the liver, we established a hemi-spleen injection mouse model of liver metastasis using Panc10.05 Cas9-mNeonGreen-expressing cells (Supplemental Figure 2E). Following transduction and selection, NT- and 230F(12)-treated cells were injected into half of the spleen of nude mice during hemi-splenectomy to seed the liver. 30 days after surgery, livers were harvested, sectioned, and stained with hematoxylin and eosin (H&E) for histology analysis (Figure 2, D and E). While we observed PC metastases in NT-treated livers (Figure 2D; black arrows), tumor regression was detected in the livers of 230F(12)-treated cells (Figure 2E; red arrows), showing that multi-targeting by CRISPR-Cas9 suppresses liver metastases.

We subsequently built a doxycycline-inducible Cas9 (Dox-iCas9) system expressing Cas9 and EGFP (doxycycline-inducible) and a U6-driven sgRNA (NT, 230F(12), and L1.4_209F). We transduced Panc10.05 with the cloned vectors, generated derivative cell lines through puromycin selection, and injected them subcutaneously into nude mice. We also prepared a no-Dox-iCas9, sgRNA-only control (“230F(12) only”), in which no EGFP would be detected in the presence of doxycycline. After tumors formed, we started doxycycline hyclate feed while measuring tumor growth (Supplemental Figure 3A). By the end of week 3, tumors were harvested, weighed, and digested for flow cytometry to detect the presence of EGFP+ cells as an indicator of PC cells (Supplemental Figure 3, B-D). Although body weights, tumor volumes, and tumor weights among treatment groups did not differ significantly (Supplemental Figure 3, A-C), we found significant decreases in PC cells in digested tumors expressing 230F(12) and L1.4_209F compared to NT control (Supplemental Figure 3D; Dunnett’s test, NT and 230F(12): *P=0.0005*, NT and L1.4_209F: *P=0.0007*, all N=3). We hypothesized that the lack of phenotypic effect was due to the incomplete induction of Cas9, as our in vitro correlates (from the same pool of cells used for xenograft) showed a 79.6% induction of Cas9-EGFP in Cas9+NT while treated with doxycycline (Supplemental Figure 3E), suggesting that ∼20% of transduced cells were not expressing Cas9. In vitro data also showed reduced growth inhibition of Cas9+230F(12), with 18.5% of living cells expressing EGFP, in contrast with prior near-complete clonogenic suppression in 230F(12)-treated cells, suggesting that reduced expression of Cas9 and/or sgRNA may be causing tumor growth. Our preliminary data suggests that multi-targeting of CRISPR-Cas9 reduces cancer cells in established tumors, and further refinement of the Dox-iCas9 model is necessary to observe a meaningful phenotypic effect.

To assess delivery to liver metastases, we used the hemi-spleen model and performed hydrodynamic injection to deliver Firefly luciferase-expressing plasmid to mouse livers (Supplemental Figure 3F). After 72 hours, livers were dissociated and sorted into two cell populations, one for liver cells (Supplemental Figure 3G) and one for PC metastatic cells based on the presence of mNeonGreen fluorescence (Supplemental Figure 3H). Intracellular staining and flow cytometry detected presence of luciferase in 23.3% of metastatic PC cells but not in liver cells (Supplemental Figure 3, G and H), demonstrating feasibility of gene therapy delivery to PC liver metastases.

### Multi-target sgRNAs exhibit on-target and minimal off-target activities

We subcultured surviving/resistant colonies from clonogenicity assays for an additional month before extracting genomic DNA to assess the targeting activity of the multi-target sgRNAs. Notably, we failed to obtain colonies from the 230F(12)- and 676F(16)-treated Panc10.05 cells, as well as from the 551R(8)-, 230F(12)-, and 164R(14)-treated TS0111 cells in all replicates (Table 1), indicating substantial cytotoxicity from these multi-target sgRNAs. We conducted whole genome sequencing (WGS) to evaluate predicted on-target and potential off-target sites (Figure 3A). Potential off-target sites with 1-4 base pair mismatches (mm) were predicted by CRISPOR (37), while off-target hits with gaps were identified using the Integrated DNA Technologies (IDT) guide RNA design checker (38). Through both manual inspection using the Integrative Genomics Viewer (IGV) and bioinformatics analyses, we found that >95% of mutations originated from on-target sites, and only 28% of 1mm sites exhibiting mutations among all potential off-targets (Table 1; Supplemental Data 1). We did not detect mutations at off-target sites containing gaps in Panc10.05 colonies (data not shown). To validate these findings, we performed targeted deep sequencing (50,000X coverage) on all potential 1mm and 2mm sites (Figure 3A), revealing mutations exclusively in 1mm sites, not in 2mm sites (Figure 3, B-C; Supplemental Figure 4, A-B). Our data demonstrate that deep NGS can reveal mutations that WGS cannot (Figure 3, B-C), likely due to the lower limit of detection by WGS (30X coverage). As an alternative approach to identify potential off-target sites, we analyzed all indels and structural variants (SVs) absent in the parental control, comparing sequence homology surrounding the mutations or breakpoints with the sgRNA sequence. None of the surrounding sequences had fewer than 5mm compared to the original sgRNA (Supplemental Table 4). We also compared SVs in non-targeted regions among individual colonies transduced with the same sgRNAs and found that these SVs were unique to each colony (data not shown). We found one shared SV instance, but its breakpoint differed from the sgRNA sequence by 13mm, suggesting it was likely present at a low level in the bulk cell line before selection by single-cell cloning.

**Figure 3.**
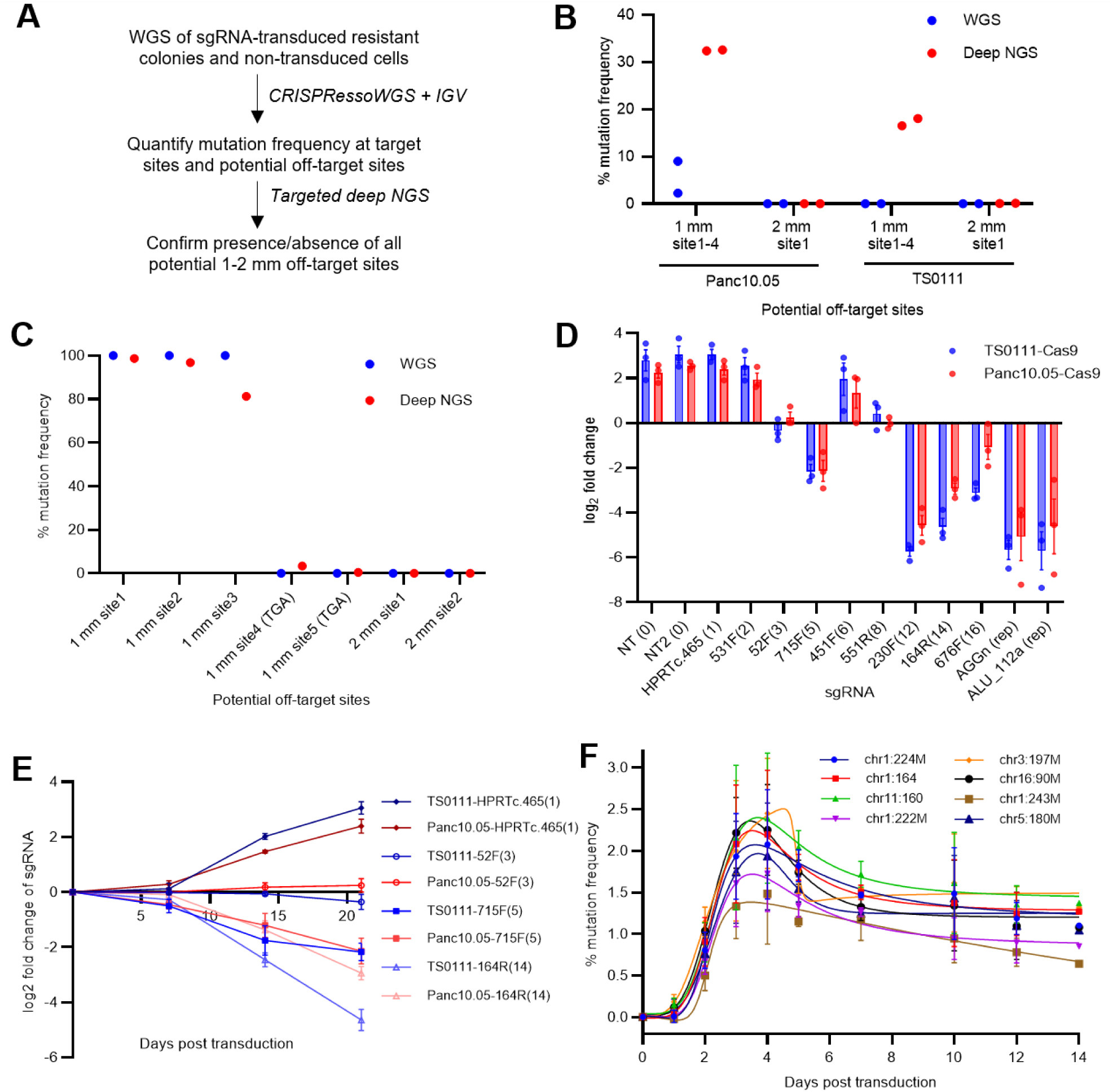
Multiple CRISPR-Cas9 scissions led to delayed cell death. **(A)** Analysis workflow for quantification of on- and off-target sites in resistant colonies from clonogenicity assays. **(B-C)** Comparisons of mutation frequency at 1-2 mismatch (mm) sites detected by whole genome sequencing (WGS) and targeted deep next generation sequencing (deep NGS) in resistant colonies. **(B)** 531F(2) sgRNA-resistant colonies. All four 1mm sites were sequenced using the same primers. N=2, mean ± SEM. **(C)** Panc10.05 164R(14) sgRNA-resistant colony. Non-canonical PAMs were indicated in parentheses. N=1. **(D)** Fold change of multi-target sgRNAs in two PC cell lines 21 days after transduction. Number of target sites in parentheses, “rep” indicates repetitive element-targeting. N=3; mean ± SEM. **(E)** sgRNA tag survival over time. N=3; mean ± SEM. **(F)** Mutation frequencies of eight 164R(14) sgRNA target sites in Panc10.05 Cas9-expressing cells at various time points. N=3; mean ± SEM. Bell-shaped least squares regression; R^2^ = 0.60-0.74. Relatively low percentages were due to the absence of antibiotic selection of transduced cells.

**Table 1.** Number of CRISPR-Cas9-induced DSBs from WGS of surviving TS0111 and Panc10.05 colonies.

| sgRNA | Number of predicted perfect target sites<sup>1</sup> | Number of potential off-target sites (1 mm - 2 mm)<sup>2</sup> | Number of mutated sites in TS0111 (0 mm - 1 mm - 2 mm)<sup>3</sup> | Number of mutated sites in Panc10.05 (0 mm - 1 mm - 2 mm)<sup>3</sup> |
| --- | --- | --- | --- | --- |
| NT | 0 | 0-1 | 0-0-0 | 0-0-0 |
| NT2 | 0 | 0-0 | 0-0-0 | 0-0-0 |
| 531F(2) | 2 | 4-1 | 2-0-0 | 2-0-0 |
| 52F(3) | 3 | 0-0 | 3-0-0 | 3-0-0 |
| 715F(5) | 5 | 2-1 | 5-1-0<sup>4</sup> | 5-1-0 |
| 451F(6) | 6 | 0-1 | 6-0-0 | 6-0-0 |
| 176R(7) | 7 | 2-1 | 6-1-0 | 6-0-0 |
| 551R(8) | 8 | 2-1 | NA | 7-0-0 |
| 230F(12) | 12 | 8-1 | NA | NA |
| 164R(14) | 14 | 5-2 | NA | 13-3-0<sup>4</sup> |
| 676F(16) | 16 | 2-6 | 16-1-0 | NA |

1. Number of perfect matches in CRISPOR using the GRCh38 human reference genome, including both canonical (NGG) and non-canonical (NGA/NAG) PAMs. 2. From CRISPOR, number of 1 and 2 mismatches (1 mm - 2 mm). 3. Matched or mismatched sites that were mutated from WGS analyses of two resistant colonies for each sgRNA, using an average variant allele frequency cutoff of 10%. Numbers are shown as 0 mm - 1 mm - 2 mm. 4. Only one colony could be obtained. NA: Resistant colonies could not be obtained.

We then compared the mutation frequencies of on-target sites across each colony to explore factors influencing variability in clonogenic inhibition (Figure 1A). Resistant colonies from 451F(6) and 176R(7) sgRNAs showed lower mutation frequencies of ∼80% and ∼36%, respectively, compared to 715F(5) (∼97%). This suggests inadequate cutting activity at the 451F(6) and 176R(7) target sites, which contributed to lower clonogenic inhibition than 715F(5) (Supplemental Figure 4C). The 451F(6) sgRNA features a TT-motif near the PAM, potentially resulting in reduced sgRNA expression when virally expressed (39). Additionally, single nucleotide variants (SNVs) were present in 4 of 6 on-target sites or PAMs of 451F(6) in Panc10.05, further affecting cutting efficiency (Supplemental Data 1). Similarly, SNVs were found in 4 of 7 on-target sites for 176R(7) in Panc10.05 and in 2 of 7 in TS0111, likely contributing to low overall mutation frequency and clonogenic inhibition (Supplemental Data 1). Furthermore, the average on-target mutation frequency of 531F(2) sgRNA-resistant colonies was higher in Panc10.05 (100%) than in TS0111 (40%; Supplemental Figure 4C), with mutation frequencies at 1mm sites higher in Panc10.05 (Figure 3B). This indicates that decreased on- and off-target activities in TS0111 contributed to the observed differences in clonogenic inhibition between the two cell lines.

We quantified the copy number (CN) of each on- and off-target site in the surviving colonies to determine the number of DSBs induced by CRISPR-Cas9. For sgRNAs lacking a corresponding surviving colony, we estimated the total number of mutated sites based on CN at all on-target sites or by assuming that the mutated sites mirrored those of a different cell line with an available colony. We found that an increased number of predicted target sites generally correlated with a higher number of mutated sites (Supplemental Figure 4D). In Panc10.05, the total copy number of target sites in the 52F(3), 715F(5), and 551R(8) colonies was comparable (9-10 cut sites), possibly explaining the similar clonogenic inhibition observed in Figure 1, A and B. The total copy number of target sites also correlated with γH2A.X foci data (Supplemental Figure 4E; Pearson r for TS0111 = 0.90, *P=0.038*; Panc10.05 = 0.98, *P=0.003*), suggesting that most observed foci resulted from CRISPR-Cas9 induced DSBs.

### Simultaneous CRISPR-Cas9 targeting leads to delayed cell death

As an independent measure of growth inhibition, we assessed sgRNA tag survival in the same cell lines, based on the premise that lethal sgRNAs would be eliminated from the sgRNA “tag” pool, while those with little or no cytotoxicity would be maintained or enriched. The 176R(7) sgRNA was excluded from subsequent analyses due to its low mutation prevalence in surviving colonies (Supplemental Figure 4C). We found that sgRNAs with a higher number of target sites exhibited greater sgRNA tag loss in Cas9-expressing cell lines, but not in parental cell lines (lacking Cas9) 21 days post-transduction (Figure 3D; Supplemental Figure 5A). We compared the sgRNA tag survival results to the results obtained from the clonogenicity assays and found that the data are significantly correlated (Spearman r for Panc10.05 = −0.78, *P = 0.001*; TS0111 = −0.92, *P < 0.0001*), indicating that sgRNA tag survival is a reliable surrogate for clonogenicity assays. We also performed sgRNA tag survival in four additional PC cell lines, revealing a general inverse correlation between the number of sgRNA target sites and sgRNA fold change, suggesting a consistent dose-response effect across six cell lines (Supplemental Figure 5B).

Interestingly, tag reduction for multi-target sgRNAs with a high number of cuts primarily occurred between days 7 and 21 post-transduction, rather than during the first 7 days (Figure 3E; Supplemental Figure 5, D and F). This was specific to Cas9-expressing cell lines, as parental cell lines showed no notable differences in sgRNA fold changes across time points (Supplemental Figure 5, A, C, and E). A comparable experiment involving the introduction of Cas9 via electroporation for transient expression yielded similar results (Supplemental Figure 5G). To investigate whether the temporal delay in tag loss was due to delayed DSB induction, we transduced the 164R(14) sgRNA into Panc10.05 Cas9-expressing cells and measured mutation frequency at 8 on-target sites over 14 days. We found that scission and repair peaked at days 3-4 (Figure 3F, R^2^ = 0.60-0.74 for all 8 sites), consistent with prior studies (40, 41). Overall, our findings indicate that DSBs induced by CRISPR-Cas9 occur within the first few days but do not immediately trigger cell death, suggesting an alternative mechanism contributes to the delay in growth inhibition.

### Ongoing and peak CIN induced by CRISPR-Cas9 scissions

To identify chromosomal changes after multiple CRISPR-Cas9 scissions, we conducted cytogenetic analyses on cells harvested from 0-21 days post-transduction of a multi-target sgRNA at 3-4 day intervals using a detailed chromosome breakage assay. The TS0111 Cas9-expressing cell line was selected for its simpler karyotype compared to Panc10.05 (Supplemental Figure 6A). We chose the 164R(14) sgRNA due to its pronounced growth inhibition in previous assays. On the first day post-transduction with 164R(14), multiple chromosome and chromatid breaks, along with radial formations, were detected (Figure 4A). Additional chromosomal aberrations accumulated over time, including ring, dicentric, and tricentric chromosomes, telomere-telomere associations, chromosome pulverizations, and endomitosis (Figure 4, A and B; Supplemental Figure 6B). Most aberrations peaked at day 14 and decreased by day 21, while chromosome/chromatid breaks remained stable throughout the experiment, indicating peak CIN during ongoing breakage events (Figure 4, A and B). We analyzed breakpoints of dicentric, tricentric, and ring chromosomes to determine their occurrence in CRISPR-Cas9 targeted versus non-targeted regions. Although SVs at targeted regions predominated at early time points, the majority of SVs occurred at non-targeted regions and peaked at day 14 (Figure 4C), consistent with ongoing chromosomal rearrangements. Notably, while most targeted regions were at telomeric regions, 61.5% of SVs at non-targeted regions were also located at telomeric sites (Supplemental Figure 6C). To investigate whether SVs at targeted regions were direct outcomes of CRISPR-Cas9 targeting, we performed a break-apart fluorescence in situ hybridization (FISH) assay at one of the 164R(14) sgRNA target sites (Figure 5, A-C). Simple rearrangements were observed at early time points (Figure 5A), while more complex rearrangements appeared later (Figure 5B). The number of cells with abnormal FISH patterns increased over time and also peaked at day 14 (Figure 5C), demonstrating ongoing chromosomal rearrangements from the initial CRISPR-Cas9 scissions.

**Figure 4.**
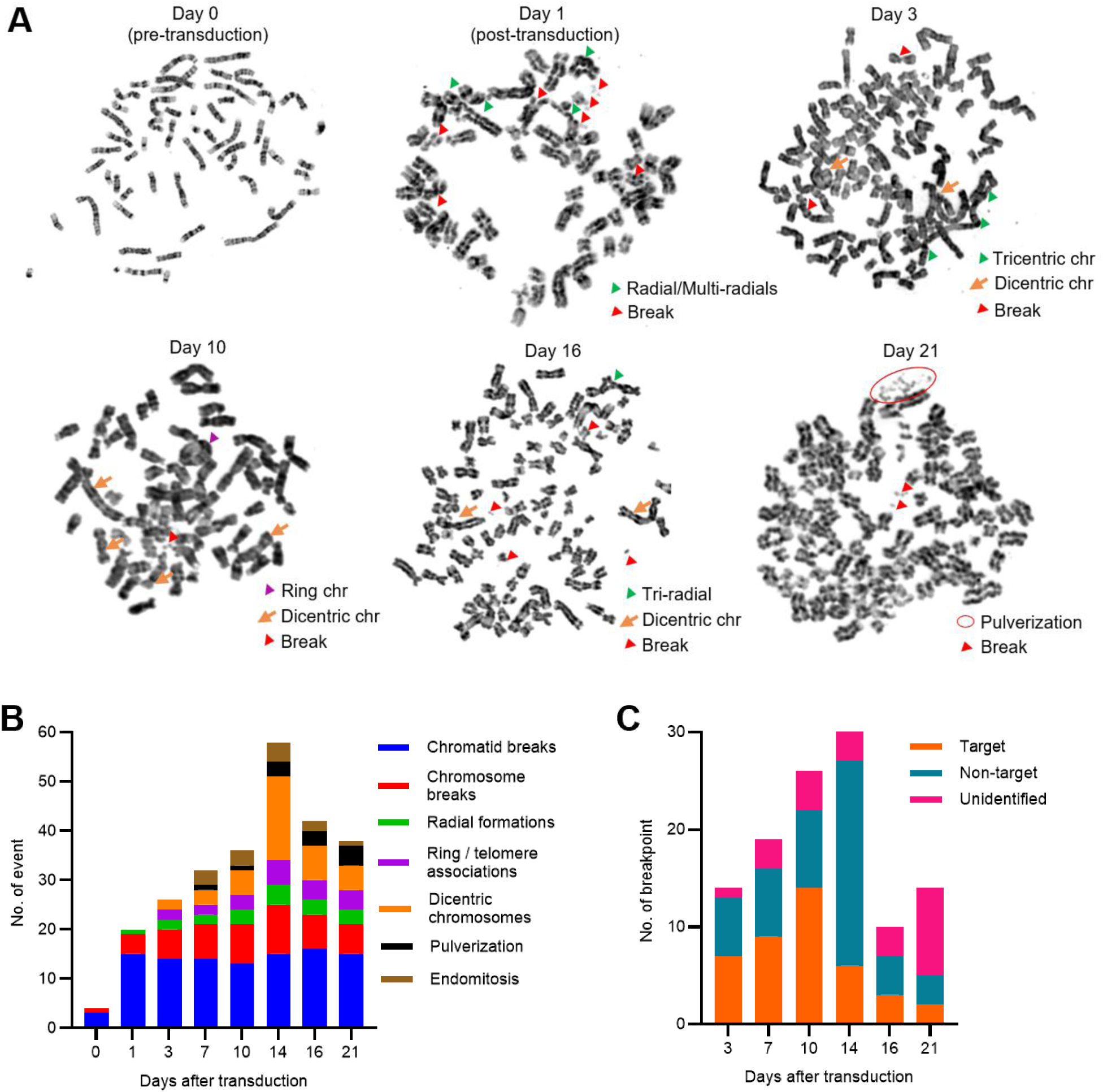
Ongoing chromosomal instability (CIN) in multi-target sgRNA transduced cells. **(A-C)** TS0111 Cas9-expressing cells were transduced with 164R(14) sgRNA and subjected to chromosome breakage assays. **(A)** Metaphase images of representative cells pre- and post-transduction of sgRNA. Karyotypic alterations are labeled. N=1. **(B)** Cytogenetic changes (events per 100 metaphase cells) over time. **(C)** Quantification of breakpoints on dicentric, tricentric, and ring chromosomes, categorized by their chromosomal band locations to determine whether the breakpoint junction was located at 164R(14) sgRNA targeted or non-targeted regions.

**Figure 5.**
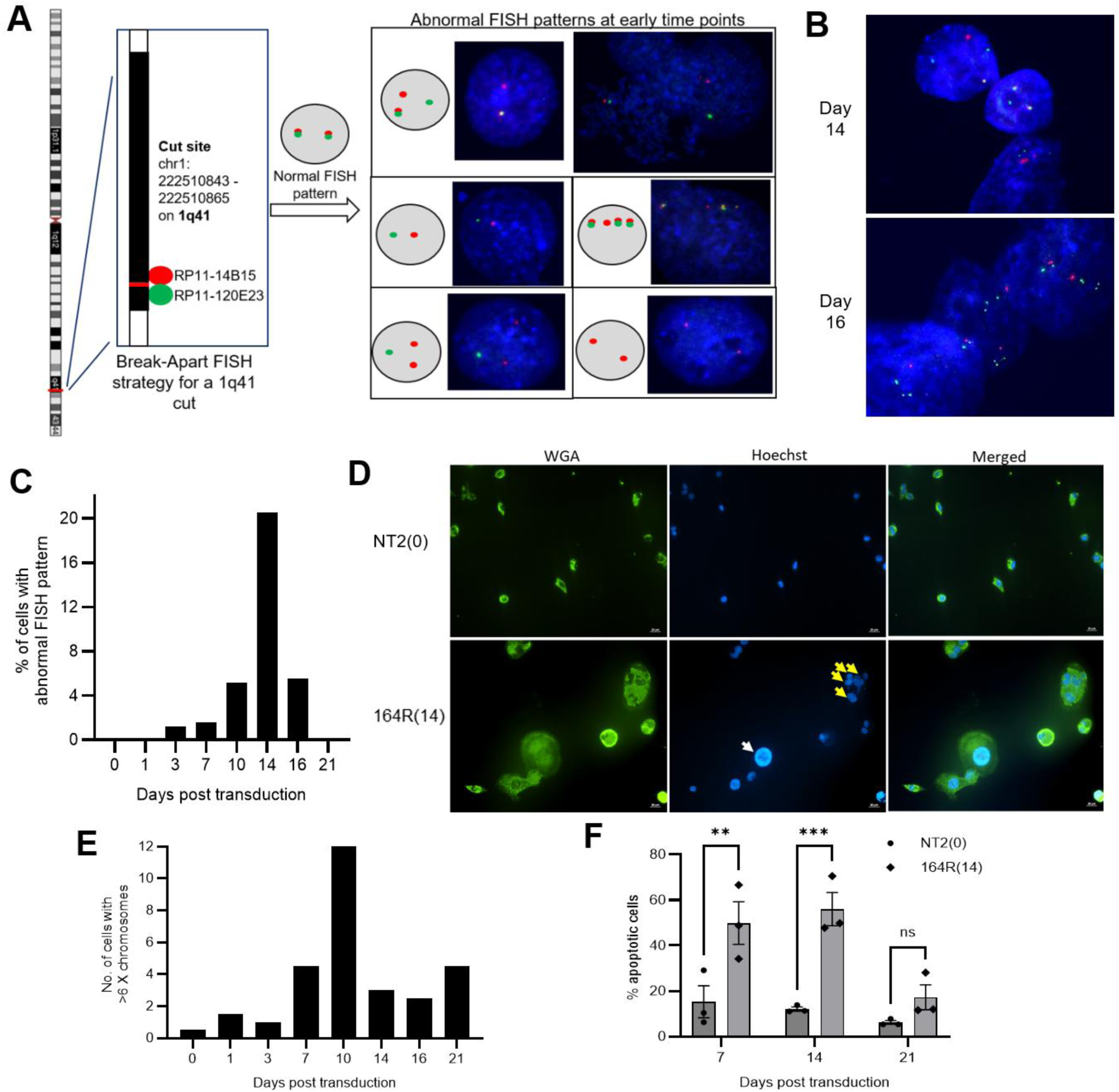
Peak polyploidy and chromosomal rearrangements in multi-target sgRNA transduced cells. **(A-C)** TS0111 Cas9-expressing cells were transduced with 164R(14) sgRNA and subjected to break-apart FISH assays. **(A)** Break-apart FISH strategy at the 1q41 cut site. Abnormal FISH patterns were shown using cells collected at early time points. DNA was stained with DAPI. N=1. **(B)** Complex rearrangements were observed in cells on 14 and 16 days after transduction. N=1. **(C)** Percentage of cells with rearrangements at 1q41 detected by break-apart FISH assay over time. **(D)** Shown are Panc10.05 cells transduced with NT2 (non-targeting) or 164R(14) and stained with wheat germ agglutinin (WGA; green) and Hoechst 33342 (blue) 14 days after transduction. White arrow: large nucleus; yellow arrows: multiple nuclei in a cell. N=3. **(E)** Number of TS0111 transduced cells with >6 X chromosomes over time using XY FISH. **(F)** Apoptosis analysis of Panc10.05 cells after treatment with 164R(14) or NT2 using Annexin V flow cytometry assay. Sidak’s multiple comparisons test, day 7: *P=0.005,* day 14: *P=0.0008,* and day 21: *P=0.53*. N=3; mean ± SEM.

TS0111 cells also exhibited polyploidy in response to the 164R(14) sgRNA, characterized by extremely large nuclei or multinucleated giant cells, with >100 chromosomes per cell detected by cytogenetic analyses (Figure 5D; Supplemental Figure 6, D and E). An X/Y FISH assay to count cells with multiple (>6) X chromosomes revealed that polyploidy peaked at day 10 and decreased by day 21 (Figure 5E). To assess if apoptosis was involved in the cell death mechanism, we analyzed apoptosis markers in cells transduced with 164R(14). The proportion of apoptotic cells stained with annexin V increased on days 7 and 14 compared to cells transduced with a non-targeting sgRNA, but decreased by day 21 (Figure 5F). We consistently encountered difficulties harvesting enough viable cells for flow cytometry analyses on day 21. However, a TUNEL (terminal deoxynucleotidyl transferase dUTP nick end labeling) assay indicated increased late-stage apoptosis on day 21 (Supplemental Figure 6F).

We also analyzed surviving colonies from our clonogenicity assays using WGS to identify, categorize, and quantify SVs absent in the parental control. This approach enabled us to examine DSB repair effects at both targeted and non-targeted regions with high resolution. Using the SV detection software manta (42), we identified SVs in surviving/resistant colonies previously transduced with multi-target sgRNAs, followed by visual inspection of all identified SVs using IGV (Figure 6). Our data revealed that while CRISPR-Cas9-induced SVs (1- and 2-target SVs) increased with the number of sgRNA target sites, the majority (87%) were non-induced DSBs (0-target SVs), which peaked in 451F(6) sgRNA-resistant colonies and subsequently decreased in 551R(8)- and 164R(14)-resistant colonies (Figure 6, A and B). Notably, translocations increased with the number of sgRNA target sites (Figure 6C). We analyzed sequences at breakpoint junctions for indels and microhomology (Figure 6, D and E), which could indicate involvement of non-homologous end joining (NHEJ), microhomology-mediated end joining (MMEJ), and single-strand annealing (SSA) (43, 44). We found that cells transduced with multi-target sgRNAs exhibited a higher proportion of breakpoints involving 1-20bp microhomologies compared to the non-targeting control (Figure 6D; Supplemental Table 5), suggesting the involvement of MMEJ in DSB repair. Overall, both cytogenetics and WGS analyses reveal that CRISPR-Cas9 scissions result in continuous chromosomal rearrangements that reach a tolerable limit, indicating that CRISPR-Cas9-induced extreme CIN is detrimental to cell survival.

**Figure 6.**
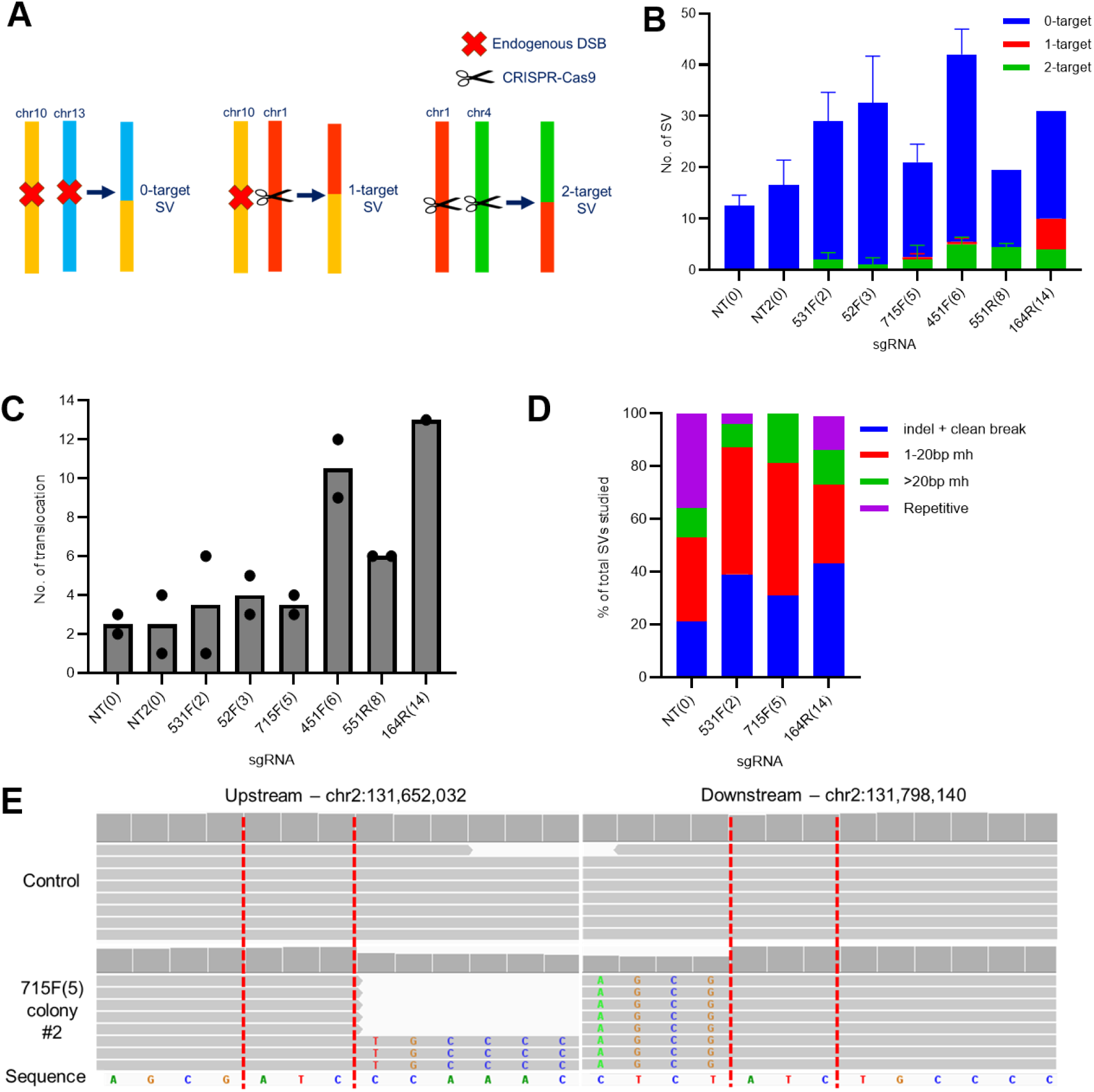
Majority of structural variants (SVs) were not directly produced from the initial CRISPR-Cas9-induced DSBs. **(A)** SVs were categorized by whether the breakpoints resulted from non-induced DSBs (0-target SV), 1 site that was CRISPR-Cas9 targeted (1-target SV), or both sites were targeted (2-target SV). **(B)** Quantification of SVs through WGS analyses of Panc10.05 surviving/resistant colonies after treatment with multi-target sgRNAs. N=2 except for 164R(14) (N=1); mean ± SEM. **(C)** Number of translocations detected in each Panc10.05 surviving colony. N=2 except for 164R(14) (N=1); mean ± SEM. **(D)** Sequences at breakpoint junctions were analyzed to identify indels and microhomology sequences (mh). Shown are the percentage of breakpoint types in each surviving colony. N=2 except for 164R(14) (N=1); mean only. **(E)** Example of a 0-target deletion from a 715F(5) sgRNA-resistant colony. The red dotted lines indicate the 3bp homology region on both upstream and downstream sequences.

### CRISPR-Cas9-induced DSBs cause higher cytotoxicity than irradiation

This CRISPR-Cas9-induced CIN resembles observations from irradiation (IR) studies, where delayed CIN and cell death manifest over several generations after IR exposure (45–47). We compared the cytotoxicity of CRISPR-Cas9-induced DSBs with IR-induced DSBs to investigate their differential effects on cancer cell growth. We first quantified DSBs induced by irradiating Panc10.05 and TS0111 with 0-10 Gy and performing γH2A.X staining, showing a clear dose-response induction of DSBs (Figure 7A; Supplemental Figure 7, A and B). Cells treated with 4Gy and higher exhibited an excessive number of foci that were uncountable (data not shown).

**Figure 7.**
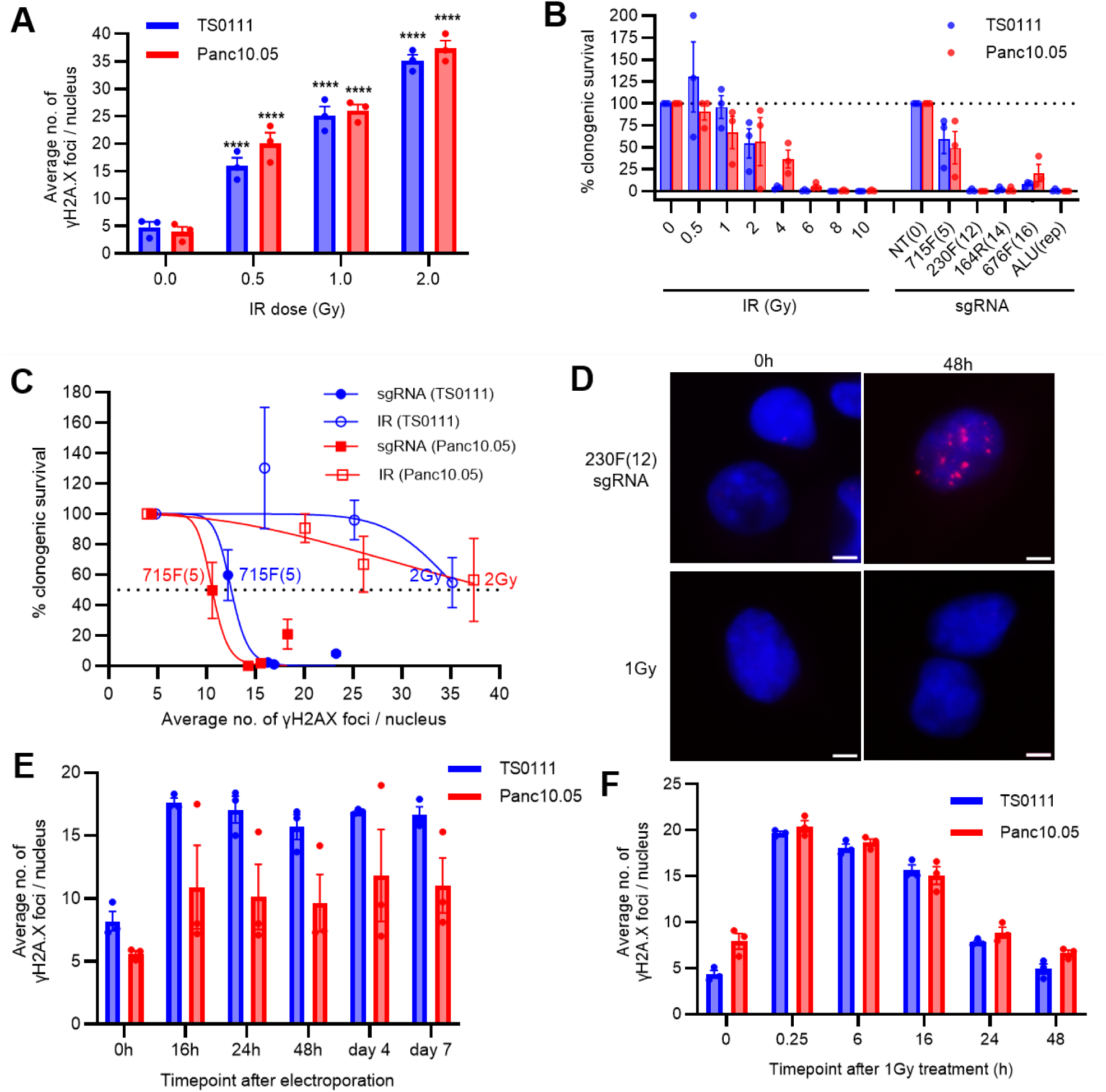
CRISPR-Cas9-induced DSBs were more cytotoxic than irradiation (IR)-induced DSBs. **(A)** Number of γH2A.X foci as a function of IR dose. >100 nuclei were analyzed for each condition. Dunnett’s test between 0Gy and each dose; **** *P<0.0001.* N=3; mean ± SEM. **(B)** Clonogenic survival as a function of IR dose or number of sgRNA target sites after 21 days. N=3; mean ± SEM, normalized to 0Gy or NT. **(C)** Clonogenic survival with increased number of γH2A.X foci detected in IR- and sgRNA-treated cells. N=3; mean ± SEM, nonlinear regressions were shown. **(D)** Representative merged images of γH2A.X (red) and DNA (DAPI, blue) staining in TS0111 cells treated with CRISPR-Cas9 RNP containing 230F(12) sgRNA or 1Gy at 0- and 48-hour timepoint. Images at 40X magnification; scale bar is 5µM. N=3. **(E-F)** Number of γH2A.X foci over time after **(E)** electroporating in CRISPR-Cas9 RNP containing 230F(12) sgRNA or **(F)** irradiated with 1Gy. >100 nuclei were analyzed for each condition. N=3; mean ± SEM.

To compare the effects of IR-induced versus CRISPR-Cas9-induced DSBs on cell survival, we assessed clonogenic inhibition in Panc10.05 and TS0111 cells following either IR or multi-target sgRNA transduction. Clonogenic inhibition increased with both IR dose and the number of sgRNA target sites 21 days post-treatment (Figure 7B). Notably, the 715F(5) sgRNA induced similar clonogenic inhibition to 2Gy IR (∼55% and ∼56%, respectively). >90% clonogenic inhibition was observed after 4Gy in TS0111 and 6Gy in Panc10.05, while >98% clonogenic inhibition was seen in 230F(12) sgRNA-treated cells from both lines. Cell viability assays supported these clonogenicity findings (Supplemental Figure 7, C and D). We also tracked cell survival in IR-treated cells every 3-4 days over 21 days, confirming that increased IR doses decreased cell numbers, and TS0111 demonstrated higher sensitivity to IR than Panc10.05 (Supplemental Figure 7, E and F).

We plotted clonogenic survival against γH2A.X foci and found that a much smaller number of DSBs induced by CRISPR-Cas9 compared to IR yielded similar levels of clonogenic inhibition (Figure 7C). For instance, while both 715F(5) sgRNA and 2Gy IR resulted in comparable reductions in clonogenic survival, the γH2A.X foci in 715F(5)-treated cells (∼11 foci) were markedly fewer than those in 2Gy-treated cells (∼36 foci). Radiation doses of 4-10Gy were highly cytotoxic to both cell lines, leading to nearly complete loss of survival, accompanied by an excessive number of foci that were uncountable. Interestingly, similar growth inhibition was observed in 230F(12)- and 164R(14)-treated cells, which had substantially fewer γH2A.X foci than those induced by equitoxic radiation doses. Our data suggests that CRISPR-Cas9-induced DSBs are considerably more cytotoxic than a comparable number of IR-induced DSBs.

Given that the cytotoxicity from CRISPR-Cas9-induced DSBs arises from ongoing chromosomal rearrangements, we hypothesized that the lower cytotoxicity of IR-induced DSBs could stem from transient DSB formation followed by immediate repair, as opposed to persistent DSBs from CRISPR-Cas9 scissions. We electroporated CRISPR-Cas9 RNP complexes containing 230F(12) sgRNAs into both PC cell lines, quantified their γH2A.X foci over time, and compared them with cells treated with 1Gy (Figure 7, D-F; Supplemental Figure 8). Since electroporated cells were not fully attached prior to the 16-hour (h) timepoint, data before 16h were excluded to avoid potential confounding effects. Our data indicated that 230F(12) sgRNA treatment led to persistent γH2A.X foci formation up to our final timepoint (day 7), with foci peaking and plateauing starting at 16h (Figure 7, D and E; Supplemental Figure 8A). High variability was observed in Panc10.05, likely due to the presence of cells that did not receive the CRISPR-Cas9 RNP complex and survived. In contrast, γH2A.X expression in irradiated cells peaked at 15 minutes and decreased over time, returning to baseline levels at 48h (Figure 7, D and F; Supplemental Figure 8B), indicating completion of DSB repair within that timeframe. To confirm that electroporation alone did not increase γH2A.X expression, we introduced a non-targeting sgRNA (NT) and counted foci at 48h, revealing an average of 5.9 foci when counting 100 cells, consistent with our baseline data. Thus, our findings suggest that while IR-induced DSBs are rapidly repaired, CRISPR-Cas9-induced DSBs persist for an extended period.

### Cells resistant to one sgRNA retain sensitivity to other sgRNAs and do not exacerbate disease

To investigate whether surviving/resistant colonies from the clonogenicity experiment (Figure 1A) enhanced tumor growth compared to non-transduced cells, we first assessed mutations at the multi-target sgRNA target sites in each colony using deep NGS, confirming their resistance against CRISPR-Cas9-induced DSBs (Figure 8A). We found that the overall mutation frequency of all target sites in each colony exceeded 99%, indicating that these colonies had their target sites mutated during the initial sgRNA transduction. We then injected both parental and resistant cells subcutaneously into nude mice and monitored tumor growth over 33 days (Figure 8B). We observed decreased tumor growth in one of the two non-targeting controls (NT #1) and in 715F(5)-resistant tumors (715F(5)), while there were no significant differences in growth for other resistant tumors compared to the parental/non-transduced control (NTC). We harvested tumors 33 days post-xenograft or postmortem (Supplemental Figure 9A) to measure their weights, and found no significant differences between resistant tumors and non-transduced controls, except for the 715F(5)-resistant tumors which displayed reduced weight (Figure 8C). No differences in body weight were noted (Figure 8D). Thus, our results indicate that DSB-resistant cancer cells induced by CRISPR-Cas9 do not grow significantly faster than the non-transduced or NT-transduced cells in subcutaneous xenograft mouse models.

**Figure 8.**
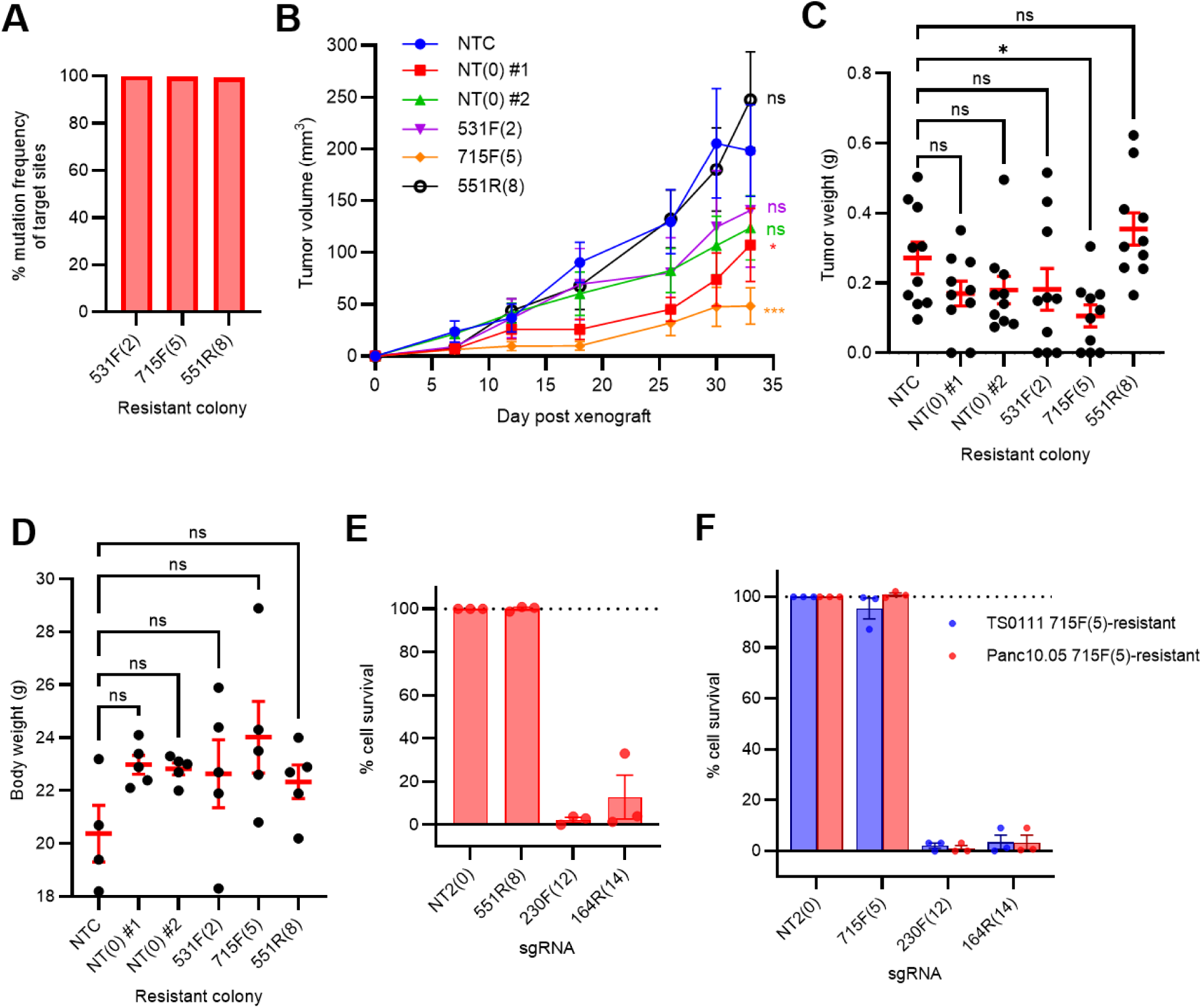
Cells resistant to one sgRNA were susceptible to other sgRNAs. **(A)** Mutation frequency of sgRNA target sites in each CRISPR-Cas9 surviving colony used for xenograft experiment. **(B-D)** Tumor growth experiment of CRISPR-Cas9 surviving colonies in subcutaneous xenograft models. In addition to a non-transduced cell line (NTC), surviving colonies from clonogenicity experiment (Figure 1A) transduced with non-targeting sgRNA (NT colony #1 and NT colony #2) and multitarget sgRNAs (531F(2), 715F(5), 551R(8)) were injected into nude mice for tumor growth. **(B)** Tumor volume measurements post-xenograft. Dunnett’s test between NTC and NT #1: *P=0.050*, NT #2: *P=0.145*, 531F(2): *P=0.349*, 715F(5): *P=0.0002*, 551R(8): *P=0.498* on week 5. N=10 (N=8 for NTC on day 30 and 33 due to early death); mean ± SEM. **(C)** Tumor weight measurements. Dunnett’s test between NTC and NT #1: *P=0.341*, NT #2: *P=0.437*, 531F(2): *P=0.457,* 715F(5): *P=0.041*, and 551R(8): *P=0.531*. N=10; mean ± SEM. **(D)** Body weight of mice 5 weeks post-xenograft. Dunn-Sidak test between NTC and the other treatment groups showed no significant differences. N=5 except for NTC (N=4 due to early death); mean ± SEM. **(E)** Cell survival of Panc10.05 551R(8)-resistant colony that was re-transduced with non-targeting sgRNA (NT2) or multi-targeting sgRNAs (551R(8), 230F(12), and 164R(14)), as detected by alamarBlue cell viability assay and normalized to NT2. N=3; mean ± SEM. **(F)** Cell survival of TS0111 and Panc10.05 715F(5)-resistant colony that was re-transduced with non-targeting sgRNA (NT2) or multi-targeting sgRNAs (715F(5), 230F(12), and 164R(14)), as detected by alamarBlue cell viability assay and normalized to NT2. N=3; mean ± SEM.

Finally, we hypothesized that cells surviving the initial CRISPR-Cas9-induced DSBs could be eliminated by a subsequent round of CRISPR-Cas9-induced DSBs. Using surviving colonies from our clonogenicity assay, with a subset of their target sites validated for mutations via NGS (Supplemental Figure 9B), we re-transduced these colonies with non-targeting sgRNA (NT2), multi-target sgRNA already present in the colonies (e.g., 715F(5)-resistant colonies were treated with 715F(5) sgRNA), 230F(12), and 164R(14) sgRNAs. After 21 days, we observed no growth inhibition in cells treated with the non-targeting control nor those re-transduced with their original sgRNAs, but found >96% inhibition with the 230F(12) sgRNA and >87% with the 164R(14) sgRNA (Figure 8, E and F; Supplemental Figure 9C). We collected double-resistant colonies (i.e., cells that survived both the original and secondary transductions) and performed NGS on their target sites. In 230F(12)-resistant colonies, the 230F(12) target sites were not mutated (Supplemental Figure 9D), suggesting that CRISPR-Cas9 activity from the secondary transduction was absent in these double-resistant colonies. We detected ∼38% mutation frequency in colonies transduced with the 164R(14) sgRNA (Supplemental Figure 9D), indicating that most surviving cells lacked CRISPR-Cas9 activity from the secondary transduction.

## Discussion

In this study, we show that simultaneous induction of multiple CRISPR-Cas9-induced DSBs overwhelms the DNA repair machinery, leading to extreme CIN and cell death - a phenomenon we term “chromosome catastrophe”. This is elucidated especially in our rare surviving colonies, which reveal that most SVs arise from this catastrophic process rather than direct CRISPR-Cas9 targeting.

Additionally, CRISPR-Cas9-induced DSBs are markedly more cytotoxic than comparable doses of IR, exhibiting persistent DSBs in sgRNA-treated cells versus transient DSBs in IR-treated cells. This is likely due to repeated DNA cleavage cycles at CRISPR-Cas9 target sites until NHEJ-mediated errors alter the sgRNA-recognition sequence, stopping further cutting (48). Such repeated cleavages have been noted in endonuclease-based systems unless the endonuclease activity is turned off (49, 50). In contrast, IR induces DSBs in a single pulse followed by immediate repair. The γH2A.X foci count in multi-target sgRNA-transduced cells may underestimate DSBs, as WGS often detects more DSBs; e.g. γH2A.X staining revealed 12 foci in TS0111 715F(5)-treated cells, while WGS detected 16 DSBs. Even accounting for background DSB formation (∼4 foci) and WGS-detected DSBs from CRISPR-Cas9, the total remains lower than that from IR. Consequently, targeting 6-8 genomic sites with CRISPR-Cas9 results in similar cancer cell kill as 2Gy IR, underscoring its therapeutic potential to induce tumor cell death similar to clinically relevant doses of radiation (51). Combining CRISPR-Cas9-induced DSBs with current chemotherapies that induce CIN, such as paclitaxel, or other agents that induce CIN selectively, such as KIF18A inhibitors, may further enhance cancer cell elimination (52–55).

Our sgRNA design strategy has led to most multi-target sgRNA sites being located near telomeres, which may contribute to the observed cell death. Cells treated with multi-target sgRNA exhibited CIN features characteristic of telomere crisis, such as extensive chromosomal rearrangements and endoreduplication, leading to a high cell death rate (56, 57). Umbreit et al. (2020) demonstrated that targeting the subtelomere of chromosome 4 in telomerase-immortalized RPE-1 cells with CRISPR-Cas9 generated chromosome bridges, supporting the hypothesis that targeting telomeric regions induces CIN (19). We observed that microhomologies were involved in most breakpoints in multi-target sgRNA-transduced colonies, aligning with literature suggesting that MMEJ plays a significant role in CRISPR-Cas9-induced DSB repair (58–60). A potential avenue for further research could involve comparing the cytotoxicity of near-telomeric versus near-centromeric targeting and exploring whether co-treatment with an MMEJ inhibitor could enhance cytotoxicity.

Polyploidization, while generally linked to therapeutic resistance (61–63), likely amplifies CIN in multi-target sgRNA-treated cells, leading to lethality rather than tumorigenicity. This could explain the absence of surviving colonies from certain treatments after three months. We observed few SVs at CRISPR-Cas9 on-target sites in surviving colonies, although CRISPR-Cas9-induced rearrangements were present in our break-apart FISH assay, suggesting these colonies may exhibit lower CIN compared to non-survivors. Future research should explore whether these colonies arise from polyploid progenies and compare their CIN rates to the bulk population.

Combining cytogenetic analysis with WGS provided complementary insights: (1) cytogenetic analysis visualizes chromosomal breakpoints directly, potentially offering a more accurate reflection than SV-calling software, which can be affected by mapping errors in repetitive regions (64); (2) WGS identifies the nucleotide sequences of breakpoints and assesses potential off-target activity at higher resolution; (3) cytogenetics reveals chromosomal aberrations contributing to CIN, while WGS relies on parameters to accurately identify structural abnormalities; (4) WGS detects smaller variations like indels and single nucleotide variants often missed by cytogenetics due to resolution limits. Thus, despite advancements in SV callers, cytogenetics remains vital for assessing genome-wide rearrangements at the single-cell level.

While we measured the copy number of individual genomic sites using WGS in the Cas9-derived line before CRISPR-Cas9 treatment, we recognize that surviving clones might exhibit unique aneuploid distributions due to genomic instability and subclonality. Our WGS analyses, conducted on two surviving clones per treatment, aimed to mitigate background variability. We acknowledge that subclones within malignant tumors can differ from the bulk cancer. In our rapid autopsy study of five patients, we identified truncal loss-of-homozygosity regions consistent across metastases, despite overall genomic instability (65).

We show that cells surviving CRISPR-Cas9 scission do not exhibit enhanced tumor growth, likely because scissions did not occur in tumor suppressor genes. Importantly, these survivors remain sensitive to further targeting, indicating resistance does not develop easily. This suggests CRISPR-Cas9 could serve as a targeted therapeutic that leverages multiple DSBs rather than modulating specific oncogenic pathways.

Successful delivery is crucial for targeting cancer in vivo. Viral delivery, commonly used in gene therapy, faces many limitations and risks, including immunogenicity, packaging capacity, and regulatory concerns (66–68). Nanoparticles, particularly lipid nanoparticles (LNPs), encapsulating Cas9 mRNA and therapeutic sgRNAs have shown promising efficacy and safety in early phase clinical trials (69–71). Although our safety data suggests well-tolerated delivery to normal cells, additional strategies may be needed to ensure preferential delivery to cancer cells (72, 73). Virus-like particles and exosome-based systems are emerging as gene therapy delivery platforms (74, 75), and more clinical data will be needed to assess their therapeutic potentials. As LNP delivery to the liver is well established and liver is a common site of metastases for various cancers (e.g. breast, colorectal, pancreatic, lung, melanoma) (76), our initial clinical trials could focus on treating liver metastases.

In summary, we demonstrate that a small number of CRISPR-Cas9-induced DSBs in non-coding regions can cause PC cell death, with a cytotoxicity more potent than IR due to accumulated CIN events leading to chromosome catastrophe. We also show that by using sgRNAs that are specific to a patient’s cancer, we could achieve tumor growth inhibition. Given the urgent need for improved therapies for PC patients, this study highlights the potential of CRISPR-Cas9 as a distinct and selective cell killing strategy against PC.

## Methods

### Mouse experiments and sex as a biological variable

Protocols for generating the Cas9-expressing cell line for xenograft and the xenograft-adapted parental line can be found in the Supplemental Methods. All mouse experiments were double-blind and used only female mice for practical reasons (e.g. less fighting).

In the subcutaneous xenograft experiment using pre-treated cells, Cas9-expressing Panc10.05 cells were transduced with lentivirus containing sgRNA-expressing plasmids at MOI 10. Following 7 days of puromycin selection, 5×10^5^ cells per tumor in 100uL PBS were subcutaneously injected into the flanks of randomized, 12-week-old female nude, athymic mice (Envigo, Indianapolis). Each mouse received two tumors, and body weight and tumor volume were measured weekly by a blinded investigator. Mice were monitored for adverse effects, and 6 weeks post-xenograft, tumors were surgically extracted and weighed by two blinded investigators.

For the hemi-spleen injection model assessing PC liver metastasis, 8-week-old female nude mice were randomized and underwent hemi-splenectomy following the pre-treatment of cells. After a week of puromycin selection, 1×10^6^ cells were injected into half of the spleen to seed the liver. Mice were anesthetized using isoflurane (2 L/min O2, 2% Isoflurane), and sterile surgical techniques were employed. Toe pinch was administered to ensure mice were at optimal anesthetic depth. Scissors and forceps were used to make a 1-cm linear incision in the dermis and peritoneal cavity and expose the spleen. Two titanium ligating clips were placed near the center of the spleen. Using scissors, the spleen was cut in half between the two ligating clips. The anterior half was placed back into the peritoneal cavity. Using a 1-mL TB syringe with a 25-gauge needle, 50uL of PBS were aspirated followed by 100uL of tumor cell suspension in DMEM tissue culture media. The final volume of 150uL was injected into the posterior half of the spleen slowly. After several seconds, two titanium ligating clips were used to clamp the vessels descending from the posterior half of the spleen before removing the posterior half. A 4-0 absorbable silk suture was used to create a continuous double stitch in the mucosa layer followed by two discrete sutures to close the skin incision. Finally, a surgical clamp was used to secure the incision further. Following surgery, mice were placed on a warming table and monitored for 30 minutes. Mice were assessed daily for discomfort. Livers were harvested 30 days after surgery, and the left lobes were fixed in 10% formalin at room temperature for 48 hours and paraffin embedded. 5um tissue sections were collected at 250-um levels between each section, yielding approximately 10 slides per liver. Tissue sections were H&E stained and evaluated by a liver pathologist.

Protocol for Dox-iCas9 mouse experiment can be found in Supplemental Methods. For the hydrodynamic injection experiment, the establishment of a hemi-spleen injection model was the same as previously described. The hydrodynamic injection protocol and luciferase protein detection were described in Supplemental Methods.

In the surviving colonies experiment, cells were maintained under blasticidin and puromycin selection for Cas9 and sgRNA expression before being injected into 6-week-old female nude mice (Envigo). Mice were randomized and received 5×10^6^ cells per tumor in 50uL matrigel (Corning) subcutaneously in both flanks. Body weight and tumor volume were measured weekly by a blinded investigator, and tumors were surgically extracted for weighing after 5 weeks post-xenograft by two blinded investigators.

### Clonogenicity and cell viability assays

For multitarget sgRNA transduction, cells were transduced with lentivirus carrying sgRNA-expressing plasmids at MOI 10 in media containing 10ug/mL polybrene. Cell culture conditions are detailed in Supplemental Methods. After 20h of incubation, cells were washed once with 1X PBS and returned in normal media. Next day, cells were diluted 1:1000 and plated in 96-well plates for clonogenic survival under 1ug/mL puromycin or 200ug/mL hygromycin selection. For electroporation of CRISPR-Cas9 RNP complex (protocol in Supplemental Methods), electroporated cells were plated evenly in 96-well plates for clonogenic growth. For normal cell line transduction, MOI was increased to 50, with 500 cells/well in 96-well plates. Negative controls included equivalent doses of NT and NT2 sgRNAs, while positive controls consisted of AGGn, L1.4_209F, and ALU_112a sgRNAs. For radiation, cells were treated with 0-10 Gy (CIXD X-RAY irradiator, xstrahl) and plated at 1:1000 and 1:10,000 dilutions on the same day. For all experiments, at the specified endpoints described in Results or figure legends, colonies were counted using phase microscopy, and alamarBlue Cell Viability Reagent (ThermoFisher) was added per the manufacturer’s instructions. Fluorescence readings were performed on a BMG POLARstar Optima microplate reader, setting excitation at 544nm and emission at 590nm, with a gain of 1000 and required value of 90%.

### γH2A.X staining and imaging

Cells were seeded at a density of 2×10^5^ cells/well of a 6-well plate with coverslips. For transduction of multitarget sgRNAs, cells were transduced and washed as described above. 48h post-transduction or at predetermined timepoints post-electroporation, cells were fixed. For the radiation experiment, cells were treated with the indicated doses of radiation (CIXD X-RAY irradiator, xstrahl) and fixed at predetermined timepoints post radiation. For both experiments, cells were fixed with 4% paraformaldehyde for 20mins at room temperature. Cells were then washed twice with 1X PBS for 5mins each, blocked and permeabilized with 5% BSA / 0.5% Triton X-100 in PBS for 30mins at room temperature, followed by an overnight incubation with anti-phospho-Histone H2A.X (Ser139) antibody (#05-636, Sigma Aldrich) at 1:1000 dilution. Next day, cells were washed thrice with PBS for 5mins each and subsequently incubated with secondary antibody conjugated with Alexa Fluor 594 (#A-11032, ThermoFisher Scientific) for 1hr at room temperature. Cells were again washed thrice with PBS and then counterstained with DAPI containing mounting medium (#H-1800, Vector Labs). Stained cells were imaged using a fluorescence microscope (Zeiss AxioImager Z1) with a 40X objective. γH2A.X foci were counted manually under the microscope with a minimum of 100 nuclei for each sample.

### WGS of surviving colonies

Genomic DNA was extracted from surviving colonies of clonogenicity assay using QIAamp UCP DNA Micro Kit (QIAGEN) according to the manufacturer’s protocol. SKCCC Experimental and Computational Genomics Core sent the samples to New York Genome Center (NYGC) for WGS with an Illumina HiSeq 2000 using the TruSeq DNA prep kit. Sequencing was carried out so as to obtain 30X coverage from 2×100bp paired-end reads. FASTQ files were aligned to both hg19 and hg38 using bwa v0.7.7 (77) to create BAM files. The default parameters were used. Picard-tools1.119 was used to add read groups as well as remove duplicate reads. GATK v3.6.0 (78) base call recalibration steps were used to create a final alignment file.

### sgRNA tag survival assay

Cells were transduced with a lentivirus pool containing all the sgRNAs in table S1 at MOI 0.1 into media containing 10ug/mL polybrene. After 24h, approximately 1 million cells were collected for day 1 timepoint, and the remaining cells were cultured for another day then subjected to 1ug/mL puromycin selection. Cells were collected on days 1, 7, 14, and 21 post-transduction and gDNA extractions were performed using QIAamp UCP DNA Micro Kit (QIAGEN) by following the manufacturer’s protocol. sgRNA library was prepared by amplifying the sgRNA target region from gDNAs using NGS primers provided by Joung et al. (79), based on the protocol outlined in the paper, and sent for NGS (Primers Table 1). Read counts of each sgRNA were extracted from FASTQ files using the script provided by Joung et al. (79) and were put through the MAGeCK (80) pipeline to obtain sgRNA fold change.

### Chromosome breakage assay

TS0111-Cas9-EGFP cells plated at 5×10^5^ /ml were treated with 164R(14) sgRNA and harvested at 0, 1, 3, 7, 10, 14, 16 and 21 days. Colcemid (0.01 μg/ml) was added 20h before harvesting. Cells were then exposed to 0.075 M KCl hypotonic solution for 30mins, fixed in 3:1 methanol:acetic acid, and stained with Leishman’s for 3 mins. For each treatment, 100 consecutive analyzable metaphases were analyzed for induction of chromosome abnormalities including chromosome/chromatid breaks and exchanges.

### Statistical analysis

The appropriate statistical tests were performed in GraphPad Prism (Version 9.2.0). The statistical models used were stated in results and figure legends. For all statistically significant results, * indicates *P<0.05*, ** indicates *P<0.01*, *** indicates *P<0.001*, and **** indicates *P<0.0001*.

### Study approval

Mouse experiments were approved by the Johns Hopkins University Animal Care and Use Committee (MO24M222). Cell line collections and uses from patient samples were approved by The Johns Hopkins Medicine Institutional Review Boards (NA_00074387).

### Data availability

The authors confirm that data supporting the findings of this study are available in the article and supplemental materials when possible. Values for all data points in graphs are reported in the Supporting Data Values file. Constructed plasmids have been deposited at Addgene, and specific sgRNA-expressing plasmids can be requested from JRE. With the exception of Panc10.05, which is an ATCC line, all other cell lines and their derivatives (including Panc10.05-derived lines) are available through Material Transfer Agreements. Sequencing data cannot be shared publicly due to IRB restrictions on de-identified data in line with research participant consent. Researchers can request more detailed data from the corresponding author through an approved collaboration arrangement.

## Supporting information

Supplemental Material, including supplemental methods, figures, and tables

## AUTHOR CONTRIBUTIONS

SSKT, AK, AB, NJR, MG, YSZ, and JRE conceived of and designed the study. SSKT, AK, AB, EHS, LM, and QZ conducted experiments. SZ, YS, AS, KB, HL, CFH, and RAA contributed to the acquisitions and analyses of data. CFH, RAA, NJR, RBS, MG, YSZ, and JRE provided resources and supervision. SSKT and JRE wrote the original draft of the manuscript. SSKT, AK, AB, EHS, LM, KB, RHH, CFH, RAA NJR, RBS, MG, YSZ, and JRE provided critical revisions to the manuscript. The authorship order among the co-first authors was determined to reflect their substantial contribution to the study, while acknowledging varying degrees of involvement in the collaborative work.

## ACKNOWLEDGEMENTS

We would like to acknowledge Drs. Michael Goggins, Hai-Quan Mao, Sarah Wheelan, Ming-Tseh Lin, Lei Zheng, Elizabeth M. Jaffee, Feyruz V. Rassool, Richard Burkhart, Christine Iacobuzio-Donahue, Jessica Gucwa, Shiwen Peng, Minh-Tam Pham, Alok Mishra, and Aparna Pallavajjala for helpful discussions. We thank Ada Tam, Raluca Yonescu, Emily Adams, Lisa Haley, Anuj Gupta, Celinia Ondeck, Jingyao Ma, Yining Zhu, Jacqueline Tang, Fidel Cai, Lee Blosser, and Victoria Stinnett for outstanding technical assistance.

## Finding

This work is the result of NIH funding, in whole or in part, and is subject to the NIH Public Access Policy. Through acceptance of this federal funding, the NIH has been given a right to make the work publicly available in PubMed Central.

National Institutes of Health grant R21CA164592 (JRE)

National Institutes of Health grant P50CA62924 (SKCCC)

National Cancer Institute CCSG P30CA006973 (SKCCC)

The Sol Goldman Pancreatic Cancer Research Center (JRE)

PanCan/AACR Innovation Award (JRE)

The STRINGER Foundation (JRE)

Casimir H. Zgonina Family Endowment for Pancreatic Cancer Research (JRE)

Dennis Troper and Susan Wojcicki (JRE)

Dick Knox/Cliff Minor Fund (JRE)

Edward A. Goldsmith Pancreatic Cancer Research Fund (JRE)

Elaine Crispen Sawyer Endowment (JRE)

Elaine T. Koehler Pancreatic Cancer Research Fund (JRE)

Eve Stancik Memorial Fund (JRE)

George Rubis Endowment for Pancreatic Cancer Research (JRE)

Hilda Buchfeller Yost Young Investigator’s Fund (JRE)

James S. McFarland Endowment for Pancreatic Cancer Research (JRE)

John J. Lussier Pancreatic Cancer Research Fund (JRE)

Linda C. Talecki Gallbladder Cancer Research Fund (JRE)

Mary M. Graf Memorial Endowment Fund for Gallbladder Cancer (JRE)

Mary Lou Wootton Endowment (JRE)

Professor J. Mayo Greenberg and Dr. Samuel L. Slovin Endowment (JRE)

Rawlings Family Pancreatic Cancer Research Fund (JRE)

## SUPPLEMENTAL MATERIAL

Supplemental Methods

Supplemental Figure 1-9

Supplemental Table 1-5

Supplemental Data 1

---

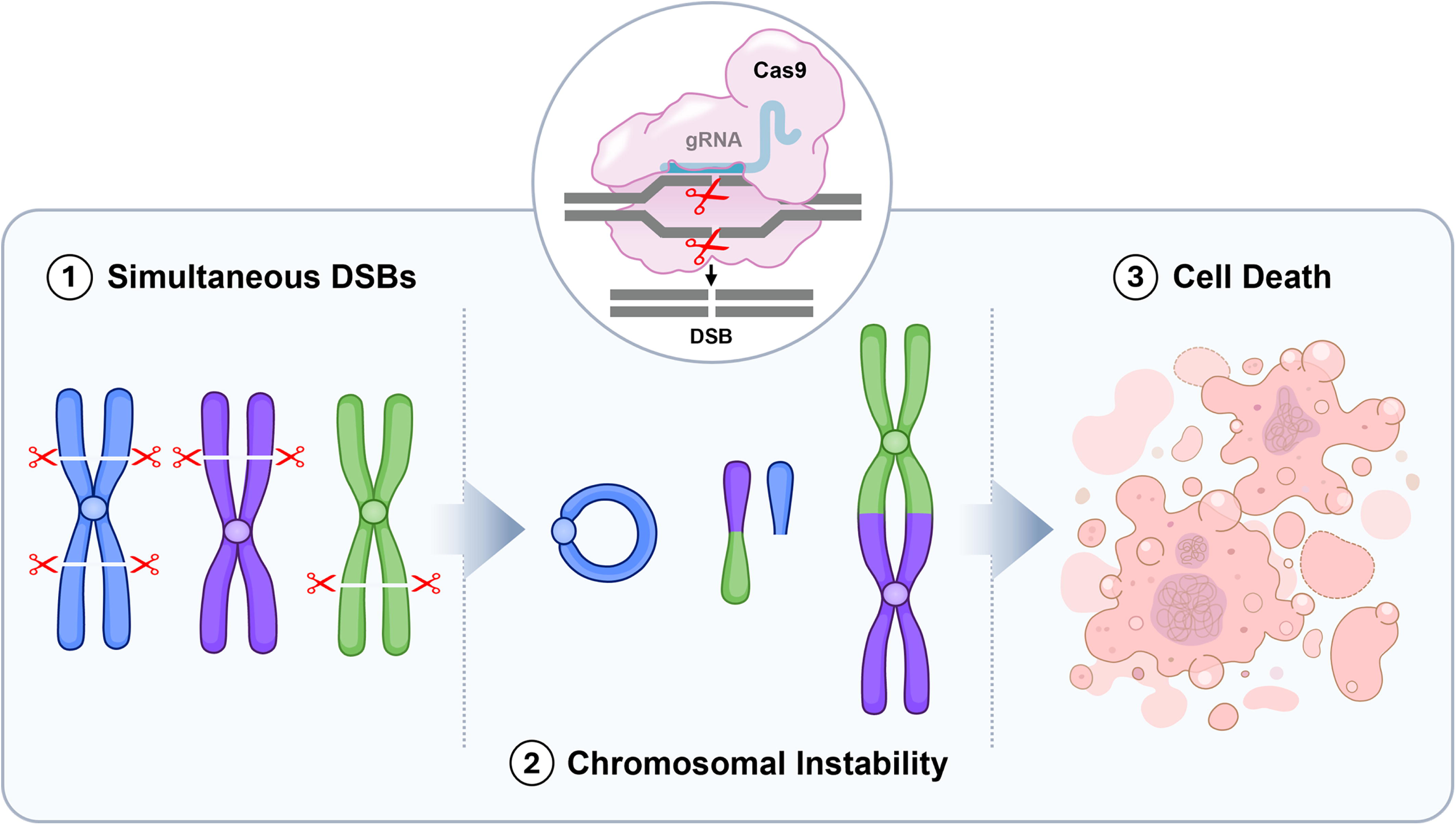

