## Supplemental Material, including supplemental methods, figures, and tables for "Simultaneous CRISPR-Cas9-induced double strand breaks are lethal in models of pancreatic cancer"

**This PDF file includes:**

Supplemental Methods

Supplemental Figure 1 to 9

Supplemental Table 1 to 5

References

**Other Supplemental Material for this manuscript includes the following:**

Supplemental Data 1

**Supplemental Methods**

*sgRNA-expressing plasmid and dox-iCas9 plasmid construction*

lentiGuide-Puro was a gift from Feng Zhang (Addgene plasmid # 52963, (1)), LentiGuide-Hygro was a gift from Caroline Goujon (Addgene plasmid # 139462, http://n2t.net/addgene:139462; RRID:Addgene_139462), and TLCV2 was a gift from Adam Karpf (Addgene plasmid # 87360; (2)). Oligonucleotides of sgRNA sequences were ordered from IDT for cloning into both lentiGuide backbones and TLCV2 backbone according to Feng Zhang's Lab Target Guide Sequence Cloning protocol (1, 3). The resulting product was transformed into One Shot Stbl3 chemically competent E. coli (ThermoFisher) according to the manufacturer's protocol and selected with both carbenicillin and ampicillin. Plasmids were extracted from ampicillin-resistant clones using QIAprep Spin Miniprep kit (QIAGEN) according to the manufacturer's protocol. Analytical digestion with restriction enzymes (NEB) was performed to verify the integrity of the plasmids, Sanger sequencing (Azenta) or whole plasmid sequencing (Plasmidsaurus) was performed to validate the insertion of sgRNA sequence.

*Cloning of Cas9-, mNeonGreen-expressing, barcoded plasmid (for xenograft)*

pLenti_Cas9_T2A_mNeonGreen_P2A_blasticidin (Addgene plasmid # 211471, (4)) was previously made. Complementary barcode oligos were ordered from IDT. The top strand oligo begins with 2bp complementary to the ClaI restriction site 5' overhang, followed by the M13F primer sequence, then followed by a unique DNA barcode of 8 bases: AATTGGCC. The bottom strand oligo begins with 4bp complementary to the XhoI restriction site 3' overhang followed by the sequence complementary to the M13F-barcode sequence of the top strand oligo.

Oligonucleotides were annealed using T4 PNK (NEB) and 10X T4 Ligation Buffer (NEB) as previously described (1). pLenti_Cas9_T2A_mNeonGreen_P2A_blasticidin vector was digested by ClaI (NEB) and XhoI (NEB) restriction enzymes for 15mins at 37°C. Digested vector was purified using the MinElute PCR Purification Kit (QIAGEN). Oligo duplexes were ligated into the digested vector using Rapid DNA Ligation Kit (ThermoFisher) according to manufacturer's instructions. The resulting product was transformed into One Shot Stbl3 chemically competent E. coli (ThermoFisher) according to the manufacturer's protocol and selected with both carbenicillin and ampicillin. Plasmids were extracted from ampicillin-resistant clones using QIAprep Spin Miniprep kit (QIAGEN) according to the manufacturer's protocol and sent for whole plasmid sequencing (Plasmidsaurus).

*Lentivirus titer preparation and quantification*

pCMV-VSV-G was a gift from Dr. Bob Weinberg (Addgene plasmid # 8454, (5)), pMDLg/pRRE and pRSV-Rev were gifts from Dr. Didier Trono (Addgene plasmid # 12251 & # 12253, (6)). 2.5ug pCMV-VSV-G, 5ug pMDLg/pRRE, 5ug pRSV-Rev, and 7.5ug transfer plasmids were used along with 50uL Invitrogen Lipofectamine 3000 reagent and 40uL P3000 reagent (ThermoFisher) for transfection into 293T cells on a 10-cm plate (95-99% confluent at transfection). Cell culture and transfection workflows were the same as the manufacturer's protocol. Upon harvesting and pooling the lentivirus-containing supernatant, the clarified supernatant was concentrated with Lenti-X Concentrator (Takara Bio) by following the manufacturer's protocol. Lenti-X qRT-PCR titration kit (Takara Bio) was used to quantify an aliquot of the clarified lentiviral supernatant according to the manufacturer's protocol.

*Cell culture*

Panc10.05 and TS0111 were kindly provided by Dr. Elizabeth Jaffee (Johns Hopkins, Baltimore, MD); A10.7 and A32.1 were gifts from Dr. Christine Iacobuzio-Donahue (Memorial Sloan Kettering Cancer Center, New York, NY); Fibro #1 and Fibro #2 were kindly provided by Dr. Ying Zou (Johns Hopkins, Baltimore, MD). All other cell lines were produced in Dr. James Eshleman's Lab (Johns Hopkins, Baltimore, MD). Panc10.05, TS0111, Panc480, Panc1002, A10.7, A32.1, Fibro #1, Fibro #2, JHH410-CAF-blast, JHH429-CAF-blast, and their derivative cell lines were STR profiled and mycoplasma tested before the start of experiments. All cells were maintained in monolayer cultures at 37˚C and 5% $CO_2$. For Fibro #1 and Fibro #2, the culture medium consisted of 1X Eagle's Minimum Essential Medium, 15% fetal bovine serum (FBS), 2mM L-glutamine, and 1X antibiotic antimycotic solution (ABAM, Sigma; contains 100u penicillin, 100ug streptomycin, and 0.25ug amphotericin B). For JHH410-CAF-blast and JHH429-CAF-blast, the culture medium consisted of 1X Dulbecco's Modified Eagle Medium (DMEM), 20% FBS, 2mM L-glutamine, and 1X ABAM. The culture medium for the remaining cell lines consisted of 1X DMEM, 10% FBS, 2mM L-glutamine, and 1X ABAM.

*Fluorescent cell line construction*

Cells were seeded at 50% confluence for 24 hours before the media was replaced to contain 10ug/mL of polybrene. Lentivirus of fluorophore-expressing plasmids were added into the media at MOI 0.01 and incubated for 18-20 hours. The media was then removed, washed once with PBS, and replaced with normal media. After 24 hours, the existing media was replaced with media that contained 5ug/mL blasticidin for a 7-day selection. BD FACSAria II or BD Fusion sorter at SKCCC Flow Cytometry Core or SKCCC High Parameter Flow Core were used for fluorescence activated cell sorting. The sorted cells were then cultured under blasticidin selection and subjected to STR profiling and mycoplasma testing. Fluorescence was verified using fluorescence microscopy before the start of experiments.

*Generation of xenograft-adapted cell lines*

For both Panc10.05 parental cells and Panc10.05-Cas9-mNeonGreen-bc6 cells, $1x10^7$ cells were injected subcutaneously into the rear flank of a female, athymic, nude mouse (Envigo) approximately 6 weeks of age. Using a 1-mL BD syringe and a 25-gauge needle, 100uL of cell suspension was delivered to the rear flank of the mouse. After 4-5 weeks, resulting tumors were explanted, finely minced, and digested with a 1:1:2 mixture of type IV collagenase (Invitrogen), hyaluronidase (Sigma-Aldrich), and 1X DMEM supplemented with 20% FBS for 1h on a shaker at 37ºC. The digested cell suspension of Panc10.05 parental cells were filtered through a 40uM cell strainer. Cells were then cultured in tissue culture flasks in regular media. For Panc10.05-Cas9-mNeonGreen-bc6 cells, the digested cell suspension was filtered through a 70uM cell strainer, resuspended in DMEM supplemented with 20% FBS, and plated on a 6-well plate coated with rat tail collagen, type I (Corning). Several days later, cells were cultured in regular media supplemented with 5ug/mL blasticidin to maintain Cas9 expression. Both cell lines were subjected to STR profiling and mycoplasma testing.

To confirm Cas9 functionality of the Panc10.05-Cas9-mNeonGreen-bc6 xenografted line, we designed a sgRNA to knock out mNeonGreen expression. Briefly, cells were plated on a 6-well plate at $1x10^5$ cells per well. Next day, cells were transduced with a sgRNA-lentivirus targeting the mNeonGreen fluorophore DNA sequence (sgRNA sequence: 5'-GGTGTGGACTTTGACATGGT-3'). Transduced cells were selected in 1ug/mL puromycin for 7 days and then examined for green fluorescence via flow cytometry. Compared to untreated cells (91.7% mNeonGreen+), those treated with an mNeonGreen-targeting sgRNA demonstrated almost no fluorescence (2.38% mNeonGreen+), indicating that Cas9 endonuclease activity was not lost during xenografting.

*Electroporation of CRISPR-Cas9 RNP complex*

Cas9 (with and without GFP), tracrRNA (with and without ATTO-488), crRNAs, and electroporation enhancer were purchased from IDT. 100uM crRNA and 100uM tracrRNA were mixed in equimolar concentrations to a final duplex concentration of 50uM, followed by heating at 95ºC for 5mins and cooling to room temperature. For Panc10.05 pool (5), the final concentration of each crRNA was 10uM instead of 50uM. Cas9 enzyme and crRNA:tracrRNA duplex components were mixed at 1:1.2 molar ratio in PBS, followed by a 20-min incubation at room temperature and on ice until electroporation (EP). Cells were resuspended in PBS ($2x10^6$ cells per EP) and combined with the RNP complex (to achieve final concentration of 2uM:2.4uM) and EP enhancer (final concentration of 2.4uM) for a total volume of 200uL per EP. Mixture was immediately transferred to pre-chilled Bio-Rad Gene Pulser 0.2cm-gap cuvettes. Electroporation parameters used on Bio-Rad Gene Pulser Xcell Electroporation Systems for TS0111: Time-constant, 250V, 2ms; and Panc10.05: Exponential, 250V, 75uF. After EP, cells were immediately transferred to pre-heated tubes with FACS buffer for cell sorting. FACS buffer was made of 1X PBS, 5% FBS, and 0.5mM EDTA. Sorting was performed on BD Fusion Sorter based on the presence of fluorophore.

*Dox-iCas9 mouse experiment*

For doxycycline-inducible Cas9 (dox-iCas9) mouse experiment, Panc10.05 cells were transduced with lentivirus containing dual dox-iCas9- and sgRNA-expressing plasmids or only sgRNA-expressing plasmids at MOI 10. Cells were puromycin selected for 7 days, then injected into female, 6-week old nude, athymic mice (Envigo). Prior to xenograft, mice were randomized by weight and housing to consist of 2-4 mice per treatment group. $1.5x10^6$ cells per tumor in 50uL matrigel (Corning) were subcutaneously implanted into the right and left flanks, with each mouse bearing two tumors. When 90% of tumors were detected 2 weeks post xenograft, Teklad Global Rodent Diet consisting of 625mg/kg doxycycline hyclate (Inotiv) was introduced into the mouse feed. Body weight and tumor volume were measured weekly by an investigator who was blinded by the treatment groups. Mice were also monitored for adverse effects. 3 weeks post xenograft, tumors were surgically extracted and weighed by two investigators who were blinded by the treatment groups.

*Hydrodynamic injection and luciferase protein detection*

Luciferase-expressing plasmid (gift from Chien-Fu Hung) was prepared at 10ug/mL in 1X PBS. Mice were administered 0.1mg/kg buprenorphine subcutaneously preceding the hydrodynamic injection. Using a 27-gauge needle and a 3-mL syringe, a volume 10% of the mouse's body weight was administered via tail vein. Mice were monitored for 5 minutes after injection until ambulatory. For luminescence imaging, mice were injected with 100uL of 500ug/mL D-luciferin (GoldBio) prior to imaging using IVIS. For antibody detection, cells were first fixed with 90% methanol for 10mins in -20˚C. After 1X PBS wash, cells were incubated in 1X PBS containing 10% normal goat serum at 4˚C for 30mins in the dark. Meanwhile, Alexa Fluor 647 Anti-Firefly Luciferase antibody (abcam; ab237252) was diluted 1/2500 in 1X PBS. After washing the cells with 1X PBS once, cells were incubated in the diluted antibody solution ($1x10^6$ cells in 100uL of solution) for 30mins at room temperature in the dark. Then, cells were washed thrice with 1X PBS and sent for flow cytometry using Attune NxT Flow Cytometer. The antibody was detected on the RL1 channel.

*Cut site determination and off-target analysis from WGS data*

WGS of surviving colonies was described in Supplementary Methods. BAM files were put into Integrated Genome Viewer (IGV (7)) to inspect all perfect and potential off-target sites (up to 4mm). Actual cut site was determined by presence of mutation (insertion, deletion, or structural variant) at the sgRNA target region. Quantification of mutation frequency of all target sites was done using CRISPRessoWGS (8). For mutations that were SVs, quantification was manually done on IGV. All predicted 1mm and 2mm sites in each colony were analyzed by PCR and deep sequencing to verify and quantify mutations (Supplementary Methods). As an alternative approach to look for off-target activity, MuTect2 v3.6.0 (9) was used to call somatic variants between the sample-control pairs. The default parameters and SnpEff (v4.1) (10) were used to annotate the passed variant calls. Manta v0.29.6 (11) was used to call somatic SVs and indels between the sample-control pairs. The default parameters were used. Variants were annotated according to UCSC refseq annotations using an in-house script. From the list of results generated, we looked for loci within the Excel files that closely matched our sgRNA sequence. This was performed with an R script that performed the following steps: 1) Read in an Excel file containing one mutation per row. 2) Obtain the forward and reverse strand sequences from the hg19 genome between the start –50bp and stop +50bp positions of the locus. 3) Align each locus's forward and reverse sequences to the target sgRNA with no gaps using the Smith-Waterman algorithm. 4) Determine the number of mismatches between the sgRNA and the nearest matching piece of DNA within each junctions. 5) Output the original information along with new columns displaying the mismatches between each junction and the sgRNA into a new Excel file. From the list of outputs, we only consider potential target sites that have <5bp mismatches to the sgRNA sequence.

*Next generation sequencing (NGS) of amplicons*

PCR was performed with primers containing partial Illumina adapter sequences to generate amplicons. Either NEBNext High-Fidelity 2X PCR Master Mix (NEB) or Platinum SuperFi II PCR Master Mix (Thermo Fisher) was used for PCR preparations, and thermocycling conditions were set based on manufacturers' suggestions. Amplicons were purified using QIAGEN MinElute PCR purification kit based on manufacturer's protocol. Purified PCR products were sent to Azenta for Amplicon-EZ service, in which 2x250bp sequencing was performed to provide ~50,000 reads per sample. FASTQ files were obtained for further analysis.

*Mutation quantification of multi-target sgRNA on- and off-target sites*

Genomic DNA from cell culture was extracted using QIAamp UCP DNA Micro Kit (QIAGEN) by following the manufacturer's protocol. Tumor genomic DNA was extracted using QIAamp DNA Mini Kit (QIAGEN) by following the manufacturer's protocol. Primers were designed using Primer3 and put through UCSC In-Silico PCR (Jim Kent) to check for primer specificity (Primers Table 2). Primers were then ordered from IDT for PCR and amplicons were sent for NGS. Mutation frequency at each site was quantified via CRISPResso2(12).

*Copy number calculation based on WGS data*

Genome-wide copy number variants from the WGS data were generated using NxClinical software version 5.2 (BioDiscovery Inc., El Segundo, CA), which was described previously (13). Briefly, two algorithms were utilized including the "Self-reference" algorithm and the "Multi-Scale Reference" algorithm. Copy number variants were detected using the hidden Markov model based on NxClinical SNP-FASST2 algorithm, with autosomal log2 ratio thresholds set at 0.7, 0.35, -0.35, and -1.5 for the detection of high-copy gains, duplications, monoallelic deletions, and biallelic deletions, respectively. Both sequencing read depths (the relative coverage) and B-allele frequencies were used to confirm copy number variant status.

*Time-course PCR*

Panc10.05-Cas9-EGFP cells were transduced with 164R(14) sgRNA and cultured without antibiotic selection. Cell pellets were collected at various time points for gDNA extraction using QIAamp UCP DNA Micro Kit (QIAGEN) by following manufacturer's protocol. Primers were designed for 8 perfect target regions of the 164R(14) sgRNA for PCR and NGS according to methods described above (Primers Table 3). Quantification of mutation frequency of all target sites was done using the CRISPResso2 (12) pipeline.

*Karyotyping*

Chromosome analyses were performed using the G-banding technique on TS0111-Cas9-EGFP cell line before and after treatment of a 14-cutter sgRNA using standard techniques. The abnormal karyotypes were described using the International System for Human Cytogenetic Nomenclature (ISCN 2020).

*1q41 Break-apart FISH assay*

FISH was performed on the TS0111-Cas9-EGFP cells before and after 164R(14) sgRNA treatment (0, 1, 3, 7, 10, 14, 16 and 21 days) using RP11-14B15 and RP11-120E23 probes flanking a 1q41 sgRNA cut according to the manufacturer's protocol (Empire Genomics Inc., Williamsville, NY). The RP11-14B15 probe binds the 5' (centromeric) side of the 1q41 sgRNA cut and in Spectrum Orange. The RP11-120E23 probe binds the 3' (telomeric) side of the 1q41 sgRNA cut and in Spectrum Green. For these probes, an overlapping red/green or fused yellow signal represents the normal pattern, and separate red and green signals indicate the presence of a rearrangement. The normal cutoff was calculated based on the scoring of the TS0111-Cas9-EGFP cells before sgRNA treatment (day 0). The normal cutoff for an analysis of 500 cells with the 1q41 break-apart probe set is calculated using the Microsoft Excel β inverse function, = BETAINV (confidence level, false-positive cells plus 1, number of cells analyzed). This formula calculates a one-sided upper confidence limit for a specified percentage proportion based on an exact computation for a binomial distribution assessment. The normal cutoff for the 1q41 break-apart probe set is 0.6% (for a 95% confidence level). For each time point, a total of 500 nuclei were visually evaluated with fluorescence microscopy using a Zeiss Axioplan 2, with MetaSystems imaging software (MetaSystems, Medford, MA), to determine percentages of abnormal cells.

*X/Y FISH assay*

FISH was performed on the TS0111-Cas9-EGFP cells before and after a 164R(14) sgRNA treatment (0, 1, 3, 7, 10, 14, 16 and 21 days) using X/Y centromere FISH probes according to the manufacturer's protocol (Abbott Molecular Inc., Des Plaines, IL). For each time point, a total of 200 nuclei were visually evaluated with fluorescence microscopy using a Zeiss Axioplan 2, with

MetaSystems imaging software (MetaSystems, Medford, MA), to determine copy number of the X chromosome. The specimen was considered abnormal if the results exceeded the laboratory-established cutoff for the X/Y probe set.

*SV identification and quantification*

From the WGS BAM files of surviving colonies, Manta v0.29.6 was used to call somatic SVs between the sample and the control, in which the control is the Panc10.05-Cas9-EGFP non-transduced cell line. The default parameters were used. Variants were annotated according to UCSC refseq annotations using an in-house script. The list of SVs generated were then individually, visually inspected on IGV to validate its presence in sample and absence in control. SVs that have passed the manual screening were quantified and characterized. Breakpoint junctions were also analyzed for microhomology manually.

*Cell membrane and genomic staining*

Alexa Fluor 488 conjugate of wheat germ agglutinin (WGA; ThermoFisher) was used to stain cell membrane on fixed cells according to the manufacturer's protocol. Hoechst 33342 stain was used to stain genomic content by incubating the cells in Hoechst 33342 for 10mins at room temperature before covering the cell with mounting media.

*Apoptosis assays*

Cells were detached using Accutase and stained with Annexin V binding antibodies and propidium iodide using BioLegend's APC Annexin V Apoptosis Detection Kit, according to the manufacturer's protocol. Fluorescence was quantified using Attune NxT Flow Cytometer. Cells were also plated on black with clear flat bottom 96-well plates and stained with both TUNEL and Hoechst using Cell Meter Live Cell TUNEL Apoptosis Assay Kit (Red Fluorescence), according to the manufacturer's protocol (AAT Bioquest). BMG POLARstar Optima microplate reader was used for fluorescence reading. For TUNEL measurement, excitation was set at 544nm and emission at 590nm, with a gain of 1000 and required value of 90%. For Hoechst 33342, excitation was set at 490nm and emission at 520nm, with a gain of 1700 and required value of 90%. Final calculation was done based on a formula used by Daniel and DeCoster (14).

**Primers Table 1-3**

Table 1. NGS primers for sgRNA tag survival assay.

| Name | Sequence* |
|---|---|
| NGS-Lib-Fwd-1 | AATGATACGGCGACCACCGAGATCTACACTCTTTCCCTACACGACGCTCTTCC GATCTTAAGTAGAGGCTTTATATATCT TGTGGAAAGGACGAAACACC |
| NGS-Lib-Fwd-2 | AATGATACGGCGACCACCGAGATCTACACTCTTTCCCTACACGACGCTCTTCC GATCTATCATGCTTAGCTTTATATATC TTGTGGAAAGGACGAAACACC |
| NGS-Lib-Fwd-3 | AATGATACGGCGACCACCGAGATCTACACTCTTTCCCTACACGACGCTCTTCC GATCTGATGCACATCTGCTTTATATAT CTTGTGGAAAGGACGAAACACC |
| NGS-Lib-Fwd-4 | AATGATACGGCGACCACCGAGATCTACACTCTTTCCCTACACGACGCTCTTCC GATCTCGATTGCTCGACGCTTTATATA TCTTGTGGAAAGGACGAAACACC |
| NGS-Lib-Fwd-5 | AATGATACGGCGACCACCGAGATCTACACTCTTTCCCTACACGACGCTCTTCCGATCTTCGATAGCAATTCGCTTTATAT ATCTTGTGGAAAGGACGAAACACC |
| NGS-Lib-KO-Rev-1 | CAAGCAGAAGACGGCATACGAGATTCGCCTTGGTGACTGGAGTTCAGACGTG TGCTCTTCCGATCTCCGACTCGGTGCC ACTTTTTCAA |
| NGS-Lib-KO-Rev-2 | CAAGCAGAAGACGGCATACGAGATATAGCGTCGTGACTGGAGTTCAGACGTG TGCTCTTCCGATCTCCGACTCGGTGCC ACTTTTTCAA |
| NGS-Lib-KO-Rev-3 | CAAGCAGAAGACGGCATACGAGATGAAGAAGTGTGACTGGAGTTCAGACGTG TGCTCTTCCGATCTCCGACTCGGTGCC ACTTTTTCAA |
| NGS-Lib-KO-Rev-4 | CAAGCAGAAGACGGCATACGAGATATTCTAGGGTGACTGGAGTTCAGACGTG TGCTCTTCCGATCTCCGACTCGGTGCC ACTTTTTCAA |

*Sequences were extracted from Joung et al Nature Protocols. 2017 Mar 23 and oligos were ordered from IDT.

Table 2. Primers for deep sequencing of surviving colonies at predicted 0-, 1-, and 2-mm sites.

| sgRNA | Chr | Up_coord | Down_coord | No_mm* | Forward primer | Reverse primer |
|---|---|---|---|---|---|---|
| 531F(2) | 1 | 366711 | 366733 | 4 1mm | GCAACTGGTCCCAAGTGTTT | AGAAGGCTGTGTGCCATCTC |
| | 9 | 98204718 | 98204740 | 2 | GGGAAGGAATTTCTCCCAAA | GGGCACAGACGAAAAGGATA |
| 715F(5) | 1 | 779526 | 779732 | 0 | GCAGAGAATTGTTTGAACCCG | GCGCATTGTAACATCAACCT |
| | 7 | 45768085 | 45768107 | 1 | CATGAGGGCTGGAAGAAAAG | CTGGGCAACAGACCAAGACT |
| | 7 | 66494590 | 66494612 | 1 | CAGCTATTCGGGAGGTTGAG | GGCTGAAGTGCAGTGGTGTA |
| | 14 | 44633068 | 44633090 | 2 | TTTCAGTTGCTTGAGATTCCA | CATCTACTCGGGAGGCTGAG |
| 551R(8) | 1 | 242992747 | 242992897 | 7 0mm | AGACGTTTCCTCACCTTCGG | AGGGCTACGCTGCTTCAATA |
| | 1 | 32466398 | 32466420 | 2 | TCTGGCTAGGACTCCCTCAA | GGGCAGACTGCTCTCAGAAG |
| 230F(12) | 10 | 133774968 | 133775123 | 7 0mm, 2 1mm | GCATTTTGGGGAGCAGAACC | GGATCCTGCCATCTGATGAA |
| 164R(14) | 11 | 159944 | 160299 | 0 | GGGATCATCACCGGACCTTT | TTTCATCATGTTGGCCAGGC |
| | 1 | 765432 | 765454 | 1 | CGGACCTTTGGCTTTTACAG | GTGCTTGGGATTATGGGTGT |
| | 10 | 38416475 | 38416497 | 1 | CACCATGCCTGCCTAATTTT | GGGTATGGGATCATCACTGG |
| | 4 | 119441239 | 119441261 | 1 | CGGACCTTTGGCTTTTACAG | CGCCTGCCTGATTTTTGTAT |
| | 7 | 66855465 | 66855487 | 1 | AAGCAGTTCTCCTGCCTCAG | GGGTGGACCTTTGGCTTTTA |
| | 7 | 66483061 | 66483083 | 1 | CCTTTGGCTTTTACAGCTTGA | TAAAGTGCCGGGATTACAGG |
| | 14 | 83660794 | 83660816 | 2 | TGCCAGTATGGCACAATGAT | TGCAGTGGCTAACCACATTAAC |
| | 2 | 112266751 | 112266773 | 2 | AAGGCAGAAATCAACAACAGA | TGTGGTTTGTGAGGATTTGG |
| 676F(16) | 16 | 90161113 | 90161265 | 13 0mm, 2 1mm | GCTTGGACAAAAGAGCCTGA | GGTTTGCAGAGCTTTCAGGG |
| | 7 | 55755509 | 55755531 | 2 2mm | TTGCTGAACACAGGAGATGC | GGTGAGAGGACACCTTGAGC |
| | 7 | 56385734 | 56385756 | 2 | GCCTCCATAAACACCCTGAA | AGCTGAGATTGTGCCACTCC |
| | 11 | 50085686 | 50085708 | 2 | GCCTCCATAAACACCCTGAA | AGAGCCAGACCCTGTCTCAA |
| | 7 | 63730827 | 63730849 | 2 | | |
| | 7 | 56846141 | 56846163 | 2 | | |

*No_mm: Number of mismatch (mm) at the CRISPR-Cas9 potential target site. “4 1mm” indicates four 1mm sites sharing the same primers.

Table 3. NGS primers for time course PCR.

| Coordinate of locus* | Primer name | Forward primer | Reverse primer |
|---|---|---|---|
| chr1:224,171,172-224,171,194 | 164R12_124M | AATGATACGGCGACCACCGAGATCTACACTCTTTCCCTACACGACGCTCTTCCGATCTTAAGTAGAGGGGATCATCACCAGACCTTTG | CAAGCAGAAGACGGCATACGAGATTCGCCTTGGTGACTGGAGTTCAGACGTGTGCTCTTCCGATCTCACCACGCCTGCCTAATTTT |
| chr1:164,976-164,998 | 164R12_164 | AATGATACGGCGACCACCGAGATCTACACTCTTTCCCTACACGACGCTCTTCCGATCTTAAGTAGAGGGGATCATCACCGGACCTTT | CAAGCAGAAGACGGCATACGAGATTCGCCTTGGTGACTGGAGTTCAGACGTGTGCTCTTCCGATCTCACCACGCCTGCCTAATTTT |
| chr11:160,165-160,187 | 164R12_1160 | AATGATACGGCGACCACCGAGATCTACACTCTTTCCCTACACGACGCTCTTCCGATCTTAAGTAGAGGGGATCATCACCGGACCTTT | CAAGCAGAAGACGGCATACGAGATTCGCCTTGGTGACTGGAGTTCAGACGTGTGCTCTTCCGATCTTTTCATCATGTTGGCCAGGC |
| chr1:222,684,185-222,684,207 | 164R12_122M | AATGATACGGCGACCACCGAGATCTACACTCTTTCCCTACACGACGCTCTTCCGATCTATCATGCTTATCACCAGACCTTCGGCTTTT | CAAGCAGAAGACGGCATACGAGATATAGCGTCGTGACTGGAGTTCAGACGTGTGCTCTTCCGATCTCACCACGCCTGCCTAATTTT |
| chr3:197,916,501-197,916,523 | 164R12_397M | AATGATACGGCGACCACCGAGATCTACACTCTTTCCCTACACGACGCTCTTCCGATCTTAAGTAGAGCACCACGCCTGCCTAATTTT | CAAGCAGAAGACGGCATACGAGATTCGCCTTGGTGACTGGAGTTCAGACGTGTGCTCTTCCGATCTGGGATCATCACCGGACCTTT |
| chr16:90,203,887-90,203,909 | 164R12_16_90M | AATGATACGGCGACCACCGAGATCTACACTCTTTCCCTACACGACGCTCTTCCGATCTTAAGTAGAGCACCACGCCTGCCTAATTTT | CAAGCAGAAGACGGCATACGAGATTCGCCTTGGTGACTGGAGTTCAGACGTGTGCTCTTCCGATCTGGGATCATCACCGGACCTTT |
| chr1:243,251,719-243,251,741 | 164R12_143M | AATGATACGGCGACCACCGAGATCTACACTCTTTCCCTACACGACGCTCTTCCGATCTATCATGCTTAGATCATCACCGGACCTTTGG | CAAGCAGAAGACGGCATACGAGATATAGCGTCGTGACTGGAGTTCAGACGTGTGCTCTTCCGATCTGCCTCAGCCTCCTAAGTAGC |
| chr5:180,721,841-180,721,863 | 164R12_580M | AATGATACGGCGACCACCGAGATCTACACTCTTTCCCTACACGACGCTCTTCCGATCTTAAGTAGAGCACCACGCCTGCCTAATTTT | CAAGCAGAAGACGGCATACGAGATTCGCCTTGGTGACTGGAGTTCAGACGTGTGCTCTTCCGATCTGGGATCATCACCGGACCTTT |

*Primers were designed for 8 of 164R(14) perfect target sites based on hg19.

### Supplemental Figures

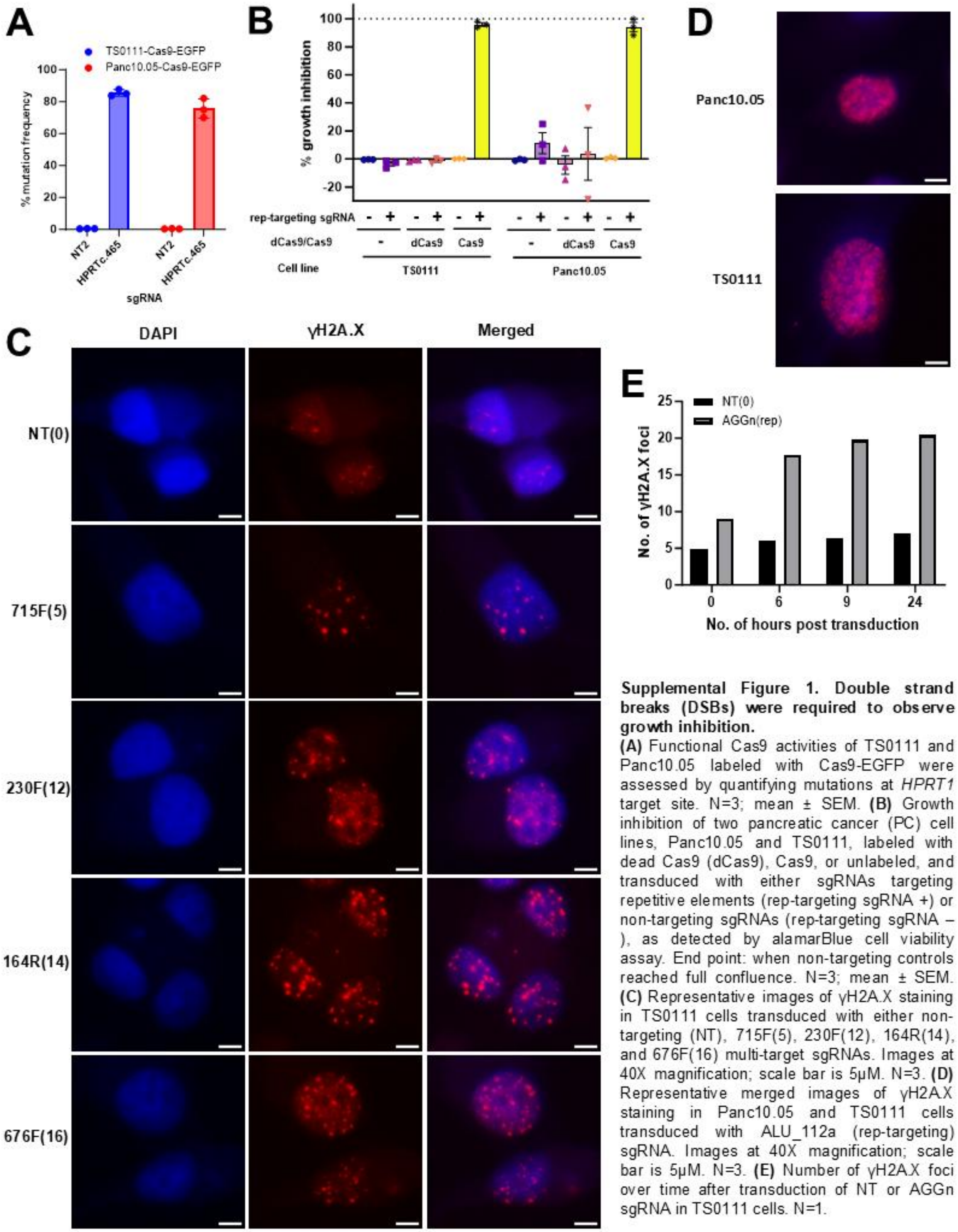


**Supplemental Figure 1. Double strand breaks (DSBs) were required to observe growth inhibition.**
**(A)** Functional Cas9 activities of TS0111 and Panc10.05 labeled with Cas9-EGFP were assessed by quantifying mutations at *HPRT1* target site. N=3; mean ± SEM. **(B)** Growth inhibition of two pancreatic cancer (PC) cell lines, Panc10.05 and TS0111, labeled with dead Cas9 (dCas9), Cas9, or unlabeled, and transduced with either sgRNAs targeting repetitive elements (rep-targeting sgRNA +) or non-targeting sgRNAs (rep-targeting sgRNA – ), as detected by alamarBlue cell viability assay. End point: when non-targeting controls reached full confluence. N=3; mean ± SEM. **(C)** Representative images of γH2A.X staining in TS0111 cells transduced with either non-targeting (NT), 715F(5), 230F(12), 164R(14), and 676F(16) multi-target sgRNAs. Images at 40X magnification; scale bar is 5μM. N=3. **(D)** Representative merged images of γH2A.X staining in Panc10.05 and TS0111 cells transduced with ALU_112a (rep-targeting) sgRNA. Images at 40X magnification; scale bar is 5μM. N=3. **(E)** Number of γH2A.X foci over time after transduction of NT or AGGn sgRNA in TS0111 cells. N=1.

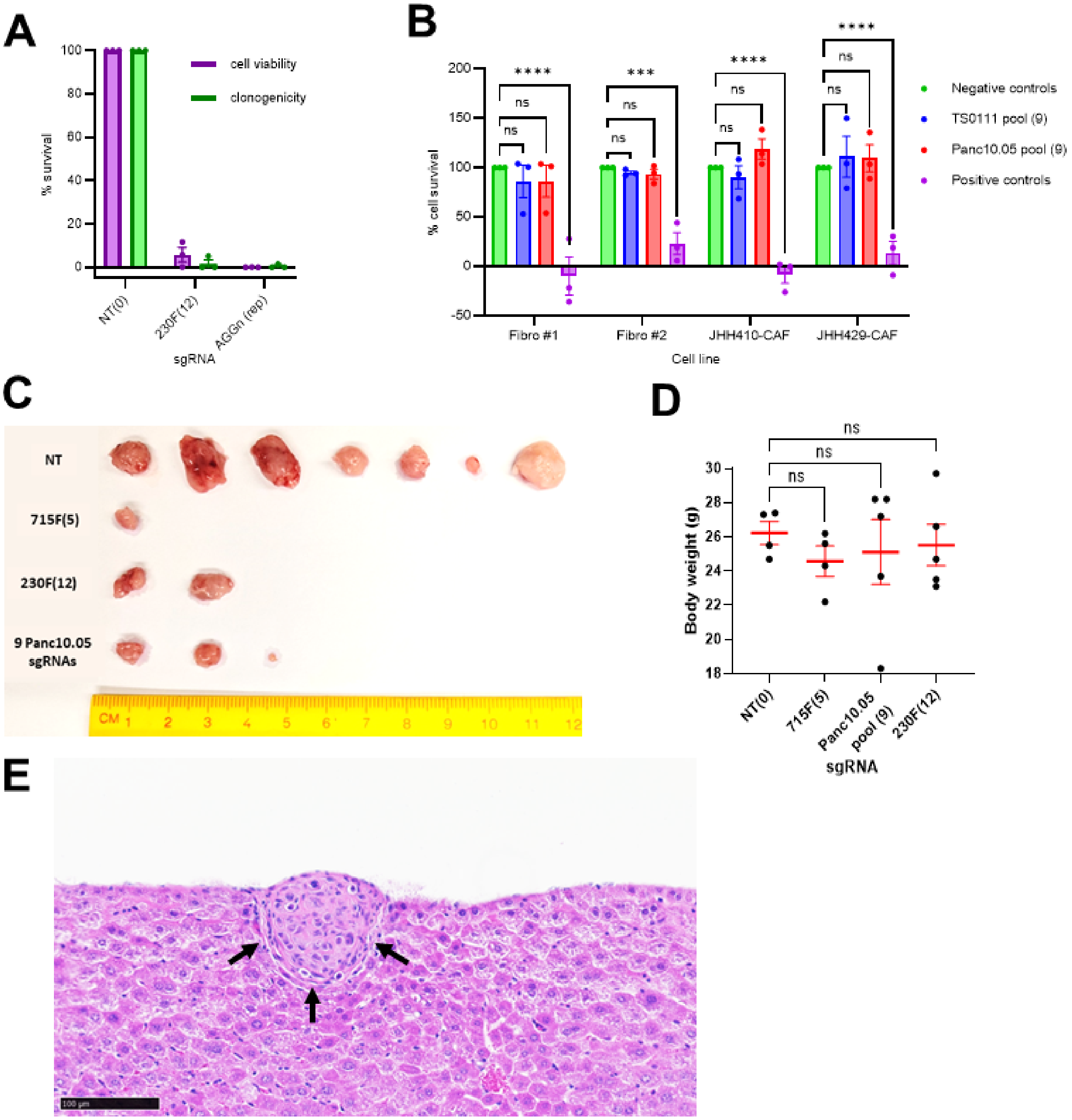


**Supplemental Figure 2. Multi-targeting by CRISPR-Cas9 inhibited tumor and metastatic growth.**
**(A)** Clonogenic and cell survival of TS0111 cells 21 days after electroporating in CRISPR-Cas9 containing multi-target sgRNAs. N=3; mean ± SEM, normalized to NT. **(B)** Cell survival of two primary skin fibroblasts lines (Fibro #1 and Fibro #2) and two cancer-associated fibroblasts lines (JHH410-CAF and JHH429-CAF) one month after the transduction of lentivirus pools containing 9 sgRNAs targeting different non-coding mutations specific to TS0111 (TS0111 pool) and Panc10.05 (Panc10.05 pool). Dunnett's test between negative controls and each treatment group; *** *P<0.001*, **** *P<0.0001*. N=3; mean ± SEM, normalized to NT. **(C)** Tumors harvested from subcutaneous mouse models on week 6 post-xenograft of pre-treated cells. **(D)** Body weight of mice 6 weeks after xenograft. Dunnett's test between NT and the other treatment group showed no significant differences. N=5 except for NT (N=4) and 715F(5) (N=4); mean ± SEM. **(E)** Hematoxylin and eosin (H&E) staining of the liver sections of a hemi-spleen injection mouse model. Black arrow: tumor cells. Images at 20X magnification; scale bar is 100μM. N=5.

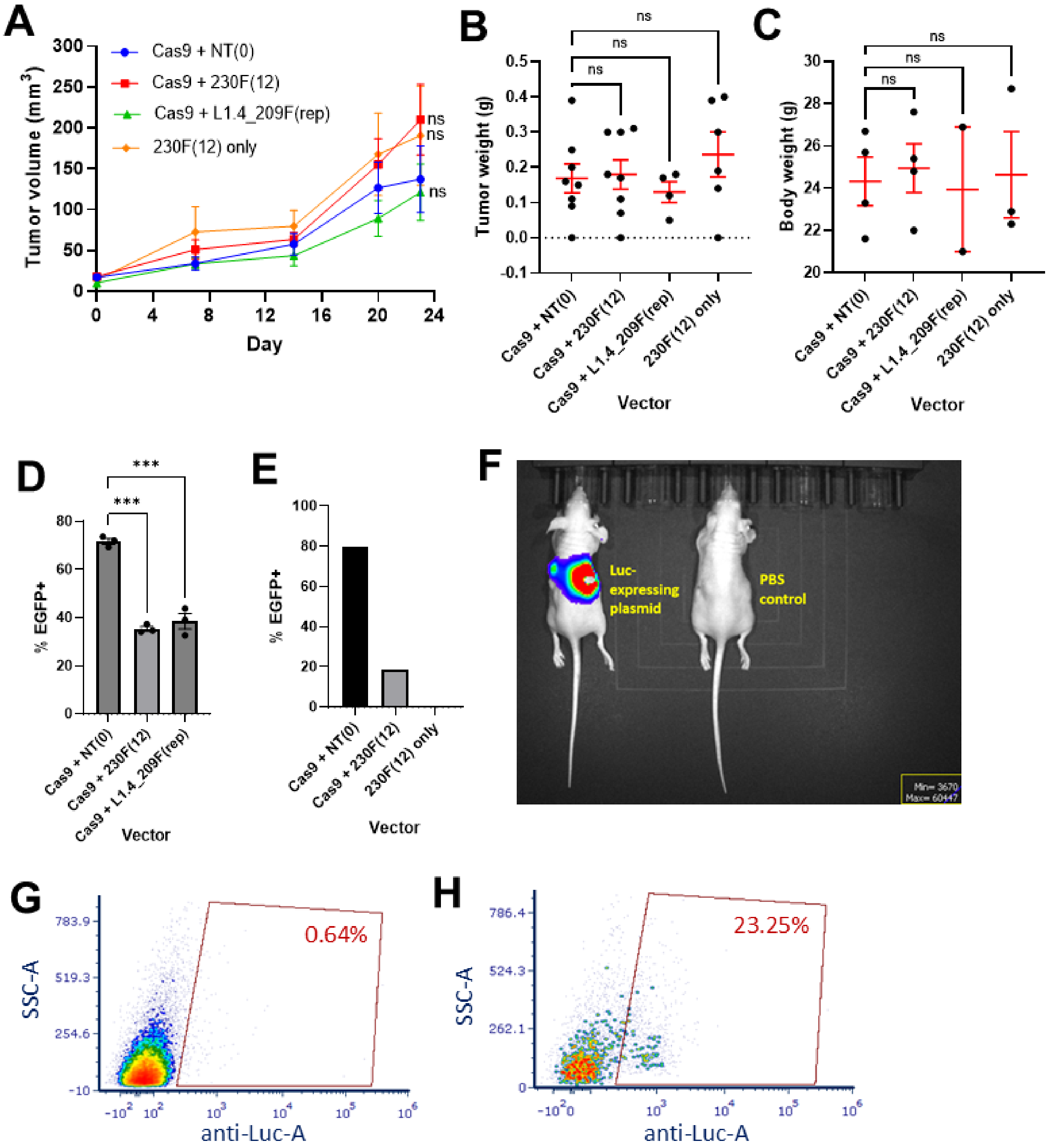


**Supplemental Figure 3. Preclinical efficacy of CRISPR-Cas9 targeting in vivo.**

**(A-E)** Dox-iCas9 mouse experiment, including measurements for **(A)** tumor volume, **(B)** tumor weight, and **(C)** body weight. Dunnett's test between NT and the other treatment groups showed no significant differences. For tumor volume and weight, N=8 except for Cas9+230F(12) (N=7) and Cas9+L1.4_209F (N=4); for body weight, N=4 except for Cas9+L1.4_209F (N=2); mean ± SEM. **(D-E)** Flow cytometry was performed on **(D)** tumors harvested along with their corresponding **(E)** in vitro correlates to detect EGFP signal (as an indicator of % of PC cells within the tumor). Dunnett's test between Cas9+NT tumors and each treatment group; *** *P<0.001*. N=3; mean ± SEM. For in vitro correlates, N=1 was shown. **(F)** IVIS image of a hemi-spleen mouse model injected with Firefly luciferase (Luc)-expressing plasmid alongside a PBS control. N=1. **(G-H)** Flow cytometry data from a digested liver 72h post-injection to identify proportion of **(G)** liver cells and **(H)** PC metastatic cells that expressed Luc. N=1.

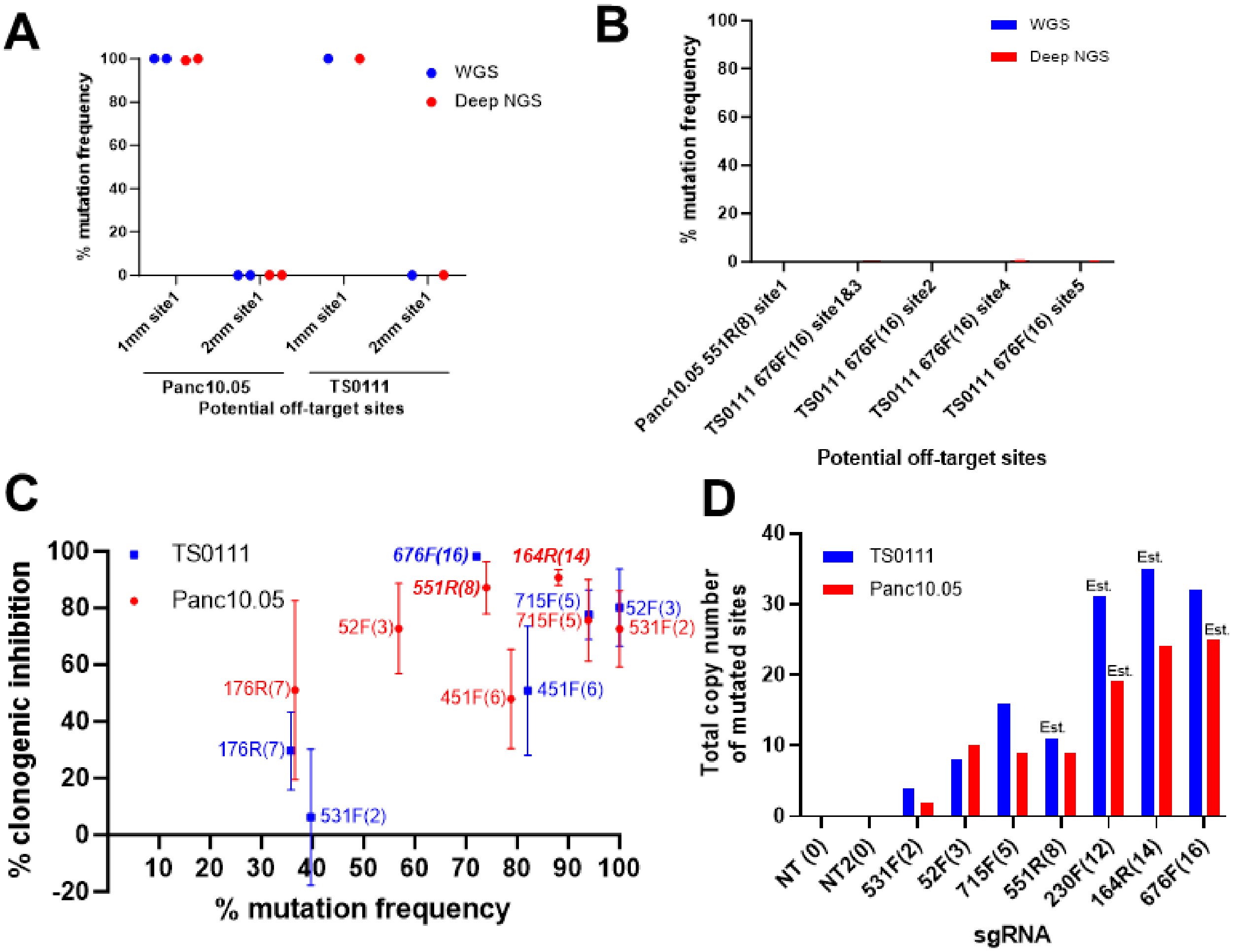


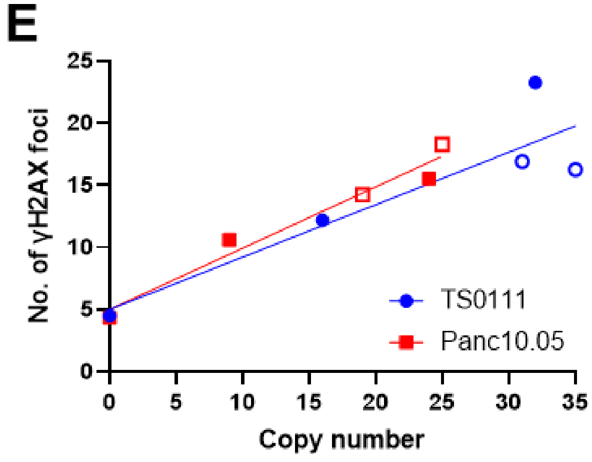


**Supplemental Figure 4. Sequencing analyses on sgRNA-resistant colonies.**
**(A-B)** Comparisons of mutation frequency at 1-2 mismatch (mm) sites detected by whole genome sequencing (WGS) and targeted deep next generation sequencing (deep NGS) in resistant colonies. **(A)** Mutation frequency analyses of 715F(5) sgRNA-resistant colonies. No amplicons detected for 1mm site2 in both cell lines. N=2 (Panc10.05) or 1 (TS0111). **(B)** Mutation frequency analyses at 2mm sites of Panc10.05 551R(8) (N=2) and TS0111 676F(16) (N=1) resistant colonies. 676F(16) 2mm site1 and site3 shared the same primers. **(C)** Average mutation frequency of all mutated on-target sites in each colony compared to clonogenic inhibition. Colonies that were present in only one cell line were bold and italicized. **(D)** Copy number of all mutated sites in surviving colonies. Est. indicates that surviving colony couldn't be obtained and the number was estimated by assuming all on-target sites were mutated or mutated sites were the same as the ones in a different cell line. **(E)** Number of γH2A.X foci as a function of the total number of mutated sites based on copy number. Emptied points indicate estimated copy number data. N=3, simple linear regressions.

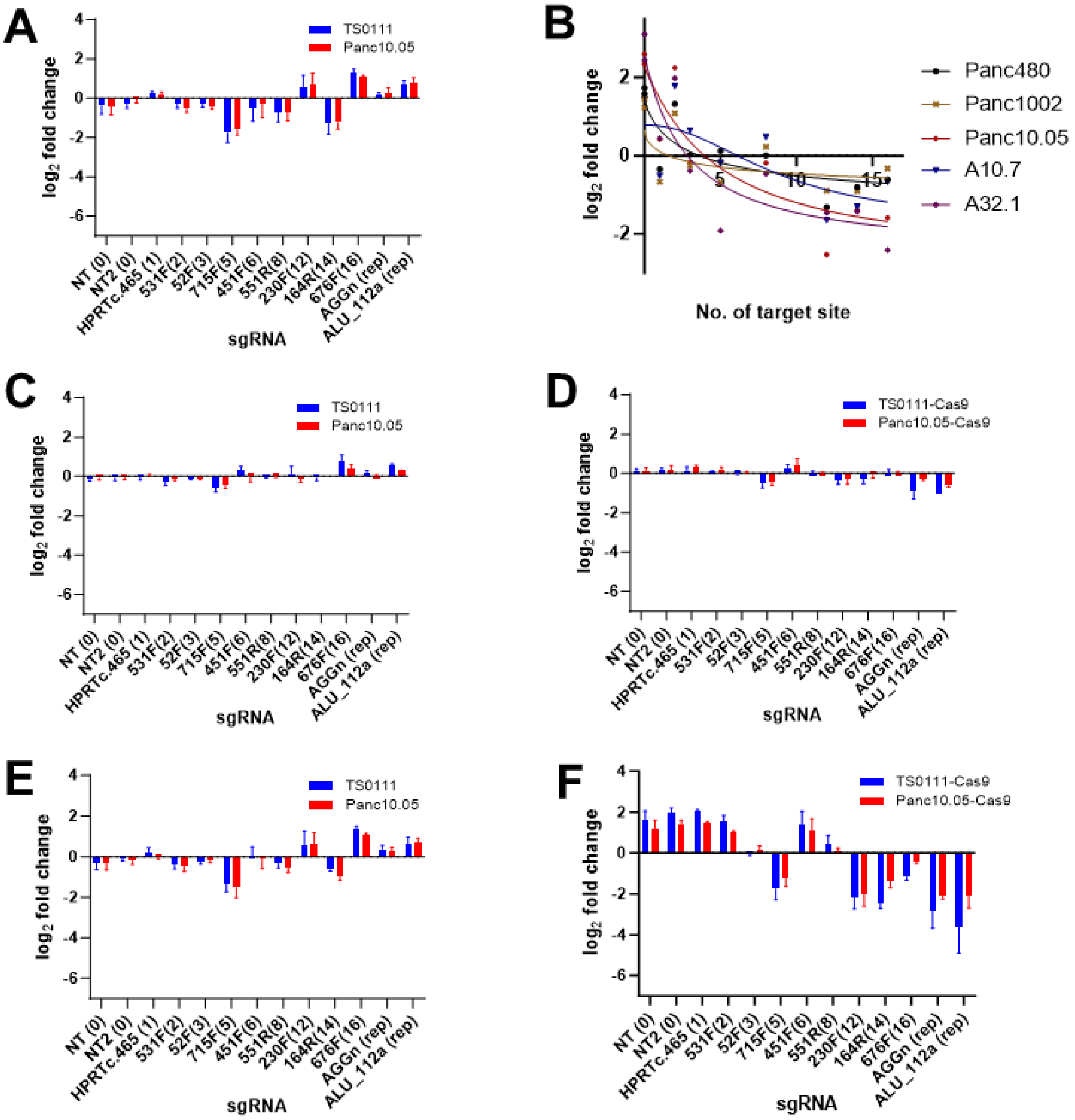


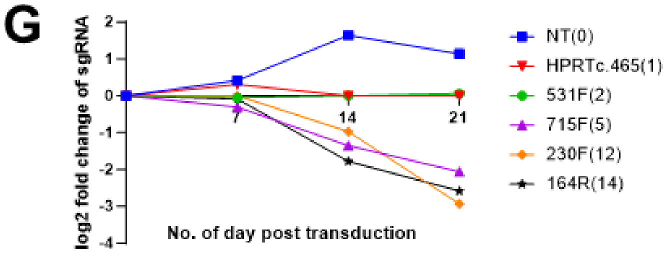


**Supplemental Figure 5. sgRNA tag loss increased over time in the presence of Cas9.** **(A)** Fold change of multi-target sgRNAs in TS0111 and Panc10.05 parental PC cell lines 21 days after transduction. N=3; mean ± SEM. **(B)** Fold change of multi-target sgRNAs in five PC cell lines transduced on day 21. N=3; mean only. **(C-D)** Fold change of multi-target sgRNAs in TS0111 and Panc10.05 **(C)** parental and **(D)** Cas9-expressing cell lines 7 days after transduction. N=3; mean ± SEM. **(E-F)** Fold change of multi-target sgRNAs in TS0111 and Panc10.05 **(E)** parental and **(F)** Cas9-expressing cell lines 14 days after transduction. N=3; mean ± SEM. Some of the data in this figure were represented in Figure 3E. **(G)** sgRNA tag survival via electroporation of Cas9 (sgRNAs were pre-introduced through lentiviral transduction) into Panc10.05 over time. N=1.

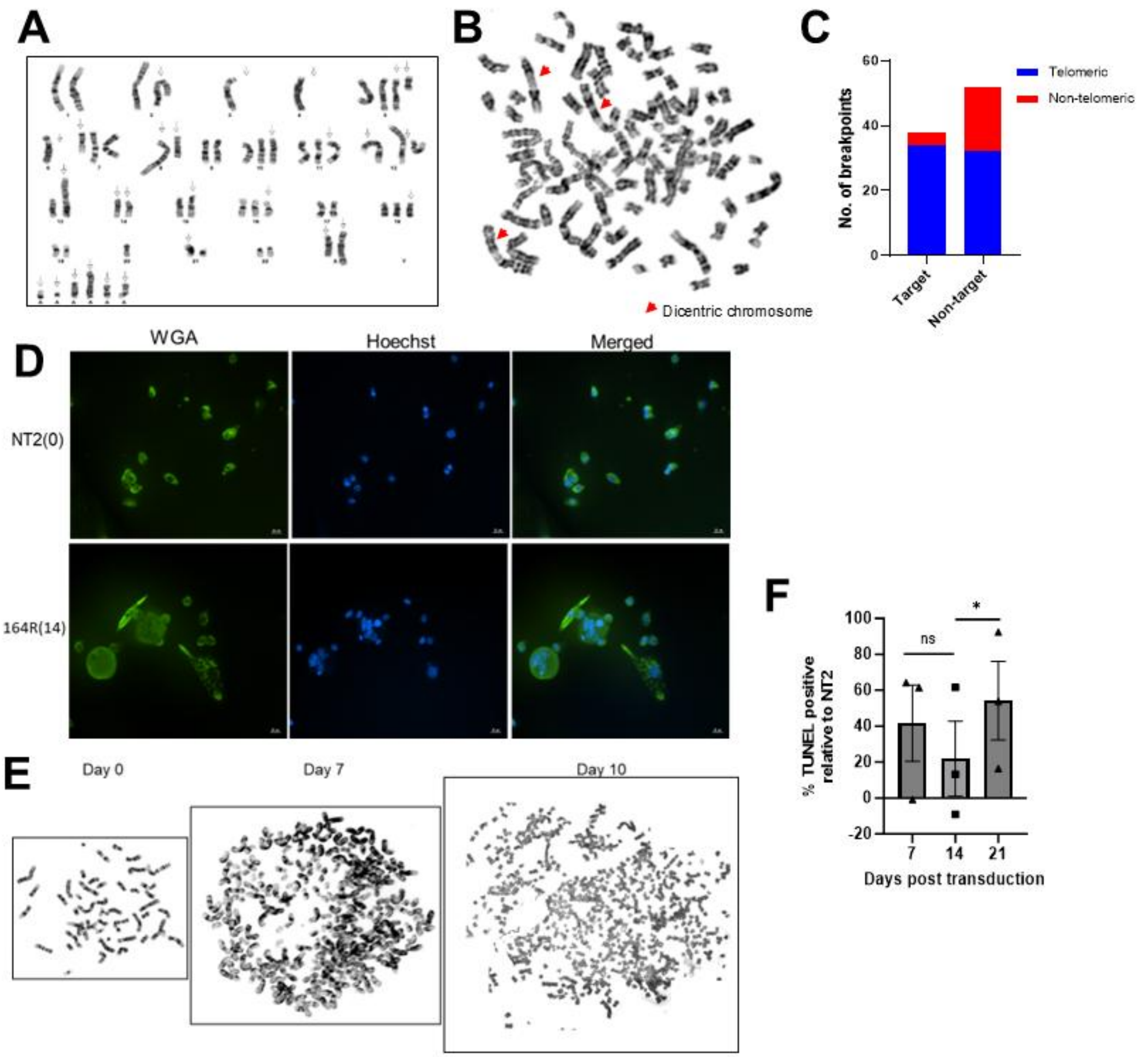


**Supplemental Figure 6. Ongoing chromosomal instability (CIN) in multi-target sgRNA-transduced cells led to cell death.** **(A)** Karyotype of TS0111-Cas9-EGFP. N=1. **(B)** Chromosome breakage analyses of 164R(14)-transduced cells on day 14. N=1. **(C)** Breakpoints of 90 dicentric and tricentric chromosomes were analyzed to determine if the breakpoint was present at a 164R(14) target region or a non-target region, and whether it was located at the telomeric end or non-telomeric regions. N=1. **(D)** Panc10.05 cells transduced with NT2 (non-targeting) or 164R(14) and stained with wheat germ agglutinin (WGA; green) and Hoechst 33342 (blue) 14 days after transduction. N=3. **(E)** Metaphase images of TS0111 cells pre-transduction (day 0), day 7 and day 10 after transduction with 164R(14). Each box contains one representative cell. N=1. **(F)** TUNEL staining to quantify Panc10.05 apoptotic cells. Paired t-test between day 7 and 14: *P=0.34*, day 14 and 21: *P=0.018*. N=3; mean ± SEM.

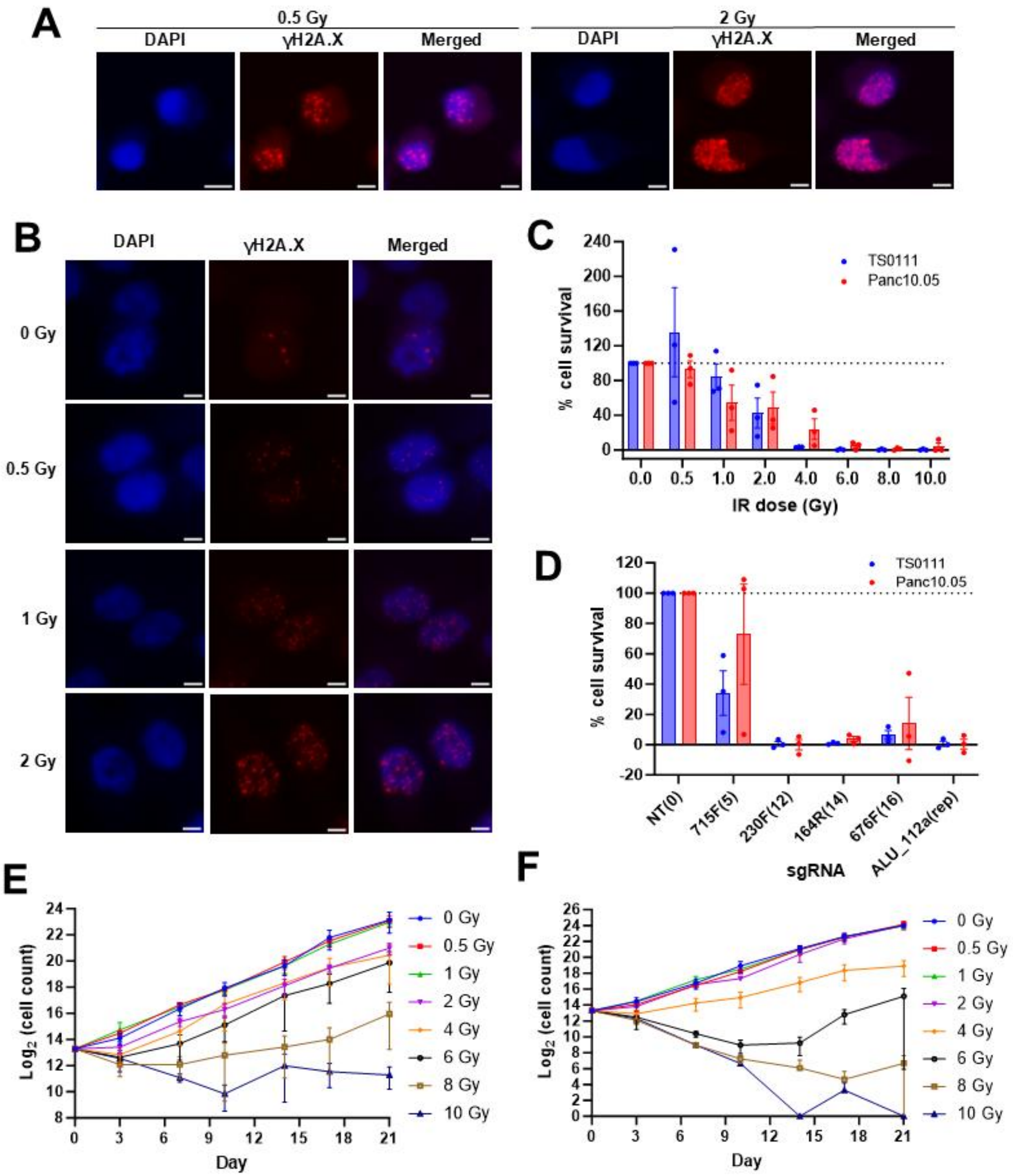


**Supplemental Figure 7. Both irradiation (IR) and multi-target sgRNAs led to growth inhibition.**
**(A)** Representative images of γH2A.X staining in TS0111 cells irradiated with 0.5 Gy and 2 Gy. Images at 40X magnification; scale bar is 5μM. N=3. **(B)** Representative images of γH2A.X staining in Panc10.05 cells radiated with 0, 0.5, 1, and 2 Gy. Images at 40X magnification; scale bar is 5μM. N=3 **(C-D)** Cell survival as a function of **(C)** IR dose and **(D)** number of CRISPR-Cas9 target sites over 21 days, as measured by alamarBlue cell viability assay and normalized to 0 Gy or NT. N=3; mean ± SEM. **(E-F)** IR-treated **(E)** TS0111 and **(F)** Panc10.05 cells were counted every 3-4 days for 21 days. N=3; mean ± SEM.

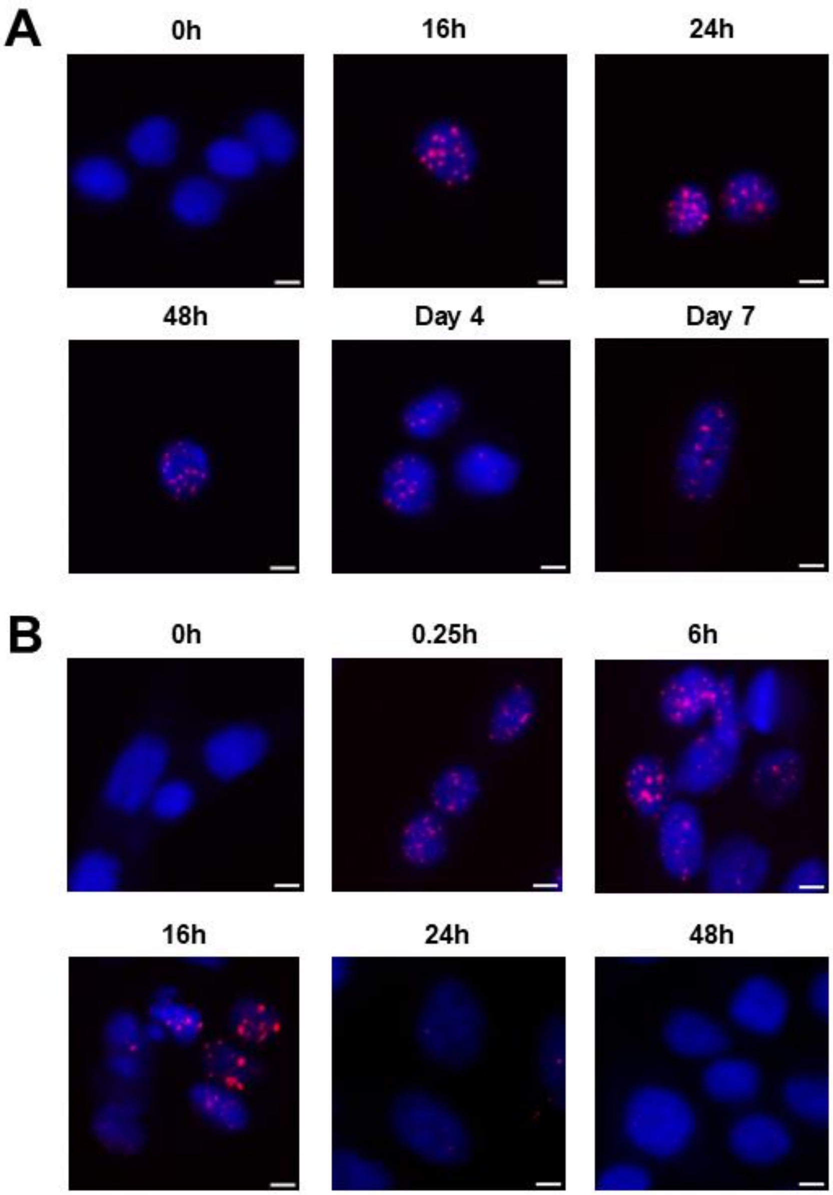


**Supplemental Figure 8. Multi-target sgRNAs caused persistent DSBs while IR triggered transient DSBs.**
Representative merged images of γH2AX staining in Panc10.05 cells treated with **(A)** CRISPR-Cas9 RNP containing 230F(12) sgRNA or **(B)** 1Gy across different timepoints. Images at 40X magnification; scale bar is 5μM. N=3.

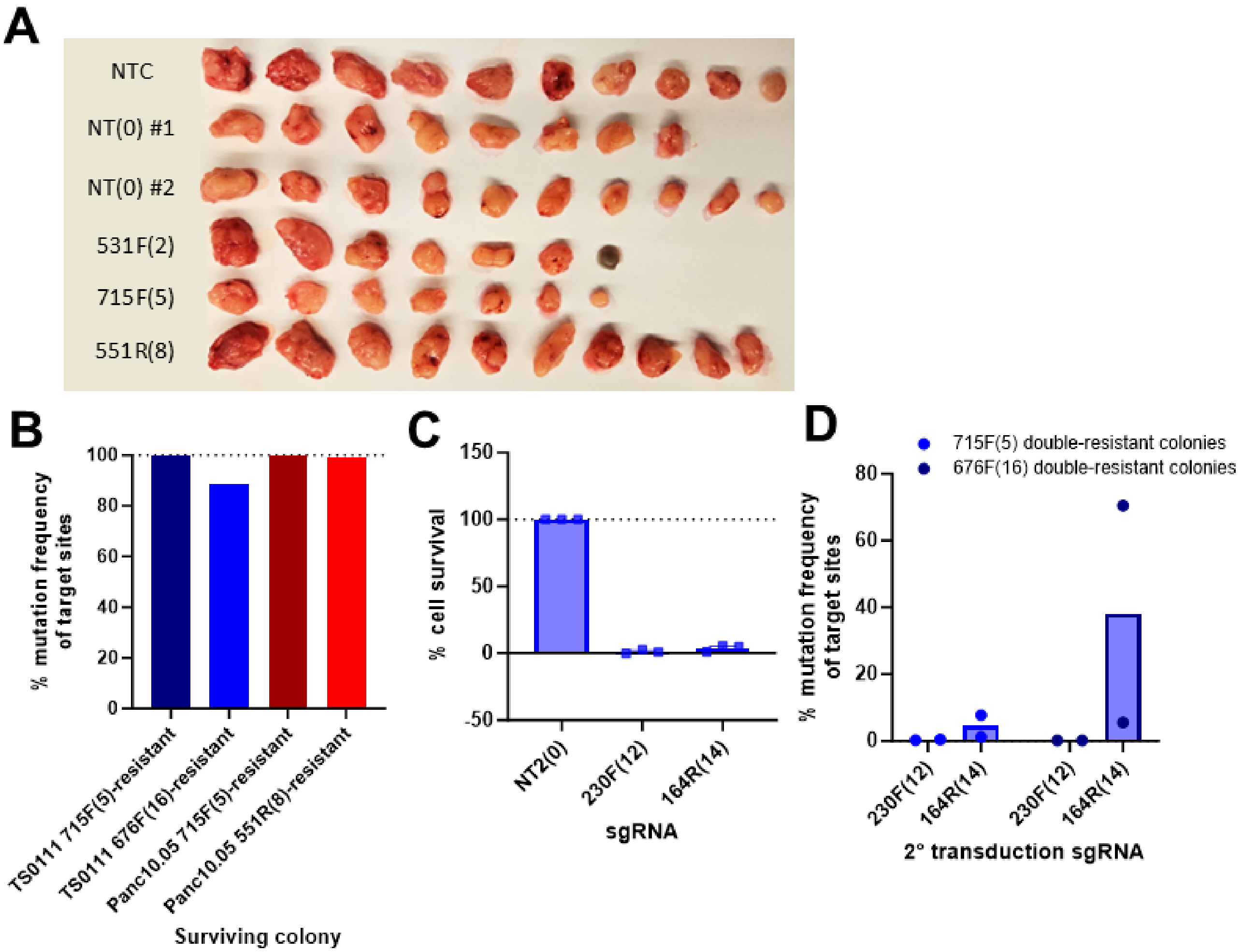


**Supplemental Figure 9. Resistance to CRISPR-Cas9 induced DSBs was resolved by a different set of DSBs.**
**(A)** Tumors harvested on day 33 post-xenograft or postmortem. The second (from the left) and last 531F(2) tumors were harvested postmortem. **(B)** Mutation frequency of sgRNA target sites in each resistant colony for secondary transduction experiment. N=1. **(C)** Cell survival of TS0111 676F(16)-resistant colony that was re-transduced with non-targeting sgRNA (NT2) or multi-targeting sgRNAs (230F(12) and 164R(14)), as detected by alamarBlue cell viability assay and normalized to NT2. N=3; mean ± SEM. **(D)** Mutation frequency of re-transduced sgRNA target sites in TS0111 715F(5)-resistant colony and 676F(16)-resistant colony. N=2; mean ± SEM.

### Supplemental Tables

#### Table 1. sgRNAs for clonogenicity and sgRNA tag survival assays.

| sgRNA | Sequence[1] | Number of perfect target sites (hg19)[2] | Number of potential off-target sites (hg19)[2] | Number of perfect target sites (GRCh38)[3] | Number of potential off-target sites (GRCh38)[3] | Doench '16 predicted efficiency score[5] | Function |
|---|---|---|---|---|---|---|---|
| NT | GTATTACTGATATTGGTGGG | 0 | 0-1-12 | 0 | 0-1-12 | NA | Negative control |
| NT2 | GCGAGGTATTCGGCTCCGCG | 0 | 0-0-2 | 0 | 0-0-2 | NA | |
| HPRTc.465 | TGGATTATACTGCCTGACCA | 1 | 0-2-8 | 1 | 0-2-8 | 64 | Functional testing |
| 531F(2) | CACTCAGCATCGACTTACGA | 2 | 4-1-0 | 2 | 4-1-0 | 66 | Experimental (Multi-target sgRNAs) |
| 52F(3) | TAATTACTGCACGATGCGCA | 3 | 0-0-2 | 3 | 0-0-2 | 59 | |
| 715F(5) | ATATATATGCGATCGAGCCC | 5 | 2-1-5 | 5 | 2-1-5 | 54 | |
| 451F(6)[4] | ACTAGTGTGCGTATGATTTG | 6 | 0-1-4 | 6 | 0-1-4 | 57 | |
| 176R(7) | TCGATGTTCTACATCGATGT | 6 | 1-1-6 | 7 | 2-1-6 | 60 | |
| 551R(8) | TTGAATTGAGTTGCAACCGA | 8 | 2-1-4 | 8 | 2-1-4 | 61 | |
| 230F(12)[4] | TTGTCCCACAATGATACTTG | 12 | 7-1-8 | 12 | 8-1-8 | 61 | |
| 164R(14)[4] | GGATATTTCACTACAGACTT | 12 | 5-2-15 | 14 | 5-2-15 | 53 | |
| 676F(16) | CTCCGAACTTAACTTGCCCT | 14 | 2-6-17 | 16 | 2-6-17 | 55 | |
| AGGn | AGGAGGAGGAGGAGGAGGAG | Repeat | | Repeat | | 37 | Positive control |
| L1.4_209F | TGCCTCACCTGGGAAGCGCA | 600 | 935-1723-2210 | 604 | 939-1710-2213 | 55 | |
| ALU_112a | TTGCCCAGGCTGGAGTGCAG | Repeat | | Repeat | | 58 | |

1. Sequences are followed in the genome by either canonical (NGG) or non-canonical (NGA/NAG) PAMs.

2 & 3. CRISPOR analysis of the sgRNAs to identify the potential on- and off-target sites (1-2-3 mismatches) in both (2) hg19/GRCh37 and (3) GRCh38 human reference genome.

4. sgRNA is labeled as inefficient by CRISPOR.

5. Cutting efficiency score based on data trained by Doench et al. 2016. Recommended for sgRNAs expressed with U6 promoter. The higher the efficiency score, the more likely is cleavage at this position.

**Table 2. Panc10.05-specific sgRNAs.**

| sgRNA name | Target Locus | Sequence *PAM* | Location type | Copy number | Potential off-target sites (0-1-2-3 mismatches) in hg38# |
|---|---|---|---|---|---|
| Panc10.05-5* | chr13:67,159,869 | GAGTGGCCTGTGATGACACT *GGG* | PCDH9 intronic | 1 | 0 - 0 - 1 - 19 |
| Panc10.05-6* | chr13:98,046,823 | GCAGAAAGAGATAGAATGGT *GGG* | intergenic | 1 | 0 - 0 - 3 - 53 |
| Panc10.05-7 | chr14:91,821,634 | CGGGTAAGAGGGAAGGATA A *GGG* | CCDC88C intronic | 1 | 0 - 0 - 3 - 44 |
| Panc10.05-8 | chr3:2,636,626 | GAAGTGAGATTCAACCATAA *TGG* | CNTN4 intronic | 1 | 0 - 0 - 0 - 14 |
| Panc10.05-10* | chr3:67,534,507 | GGGCCTTACCTGAAAGCAGC *AGG* | SUCLG2 intronic | 1 | 0 - 0 - 1 - 23 |
| Panc10.05-11* | chr3:76,973,237 | AGAATTTGAGCGACAGTATG *TGG* | ROBO2 intronic | 1 | 0 - 0 - 2 - 9 |
| Panc10.05-12* | chr4:133,652,679 | TAACACCCTTCTCTGAACAA *TGG* | intergenic | 1 | 0 - 0 - 1 - 29 |
| Panc10.05-14 | chrX:65,772,386 | GGACCTGGACTTCTTCAAGG *AGG* | intergenic | 2 | 0 - 0 - 2 - 26 |
| Panc10.05-15 | chrX:126,458,293 | TGACGTCATCTGTAATATTG *CGG* | intergenic | 2 | 0 - 0 - 2 - 29 |

*sgRNAs used for electroporation.

#CRISPOR analysis of the sgRNAs to identify the potential perfect and off-target sites (1-2-3 mismatches) in the hg38 human reference genome.

**Table 3. TS0111-specific sgRNAs for immunofluorescence (IF) staining and co-culture assays.**

| sgRNA name | Target | sgRNA sequence | PAM | Location type | Potential off-target sites (0-1-2-3 mismatches) in hg38* |
|---|---|---|---|---|---|
| TS0111-2 | chr10:70092203 | AATTCAGTGGACGACGCCGA | GGG | PBLD intronic | 0 - 0 - 0 - 1 |
| TS0111-3 | chr12:10876057 | TCATTAGCATTTAAAGGCGC | CGG | Intergenic | 0 - 0 - 0 - 8 |
| TS0111-7 | chr12:106150227 | AATTAGCCGGAGTGGTGGTG | GGG | Intergenic | 0 - 0 - 33 - 48 |
| TS0111-8 | chr12:128055569 | ACATGGTGCCCCGTCGGCTA | CGG | Intergenic | 0 - 0 - 0 - 2 |
| TS0111-9 | chr12:130765849 | TGGGCCCAGGCTCGGGGGCT | GGG | Intergenic | 0 - 0 - 7 - 50 |
| TS0111-11 | chr14:30809300 | AATCATGATGTCTGTCTTCA | TGG | Intergenic | 0 - 0 - 3 - 63 |
| TS0111-12 | chr14:39901580 | CAGCGGCCCGGAAGCCTCAA | GGG | FBXO33 UTR | 0 - 0 - 0 - 5 |
| TS0111-17 | chr14:91885563 | TTAATTGCTTCTCCGCCCGC | CGG | Intergenic | 0 - 0 - 0 - 1 |
| TS0111-18 | chr16:14136787 | GGCTTTGTTTATGGGACAGA | TGG | Intergenic | 0 - 0 - 3 - 21 |

*CRISPOR analysis of the sgRNAs to identify the potential perfect and off-target sites (1-2-3 mismatches) in the hg38 human reference genome.

**Table 4. Lowest number of mismatch (mm) of mutations detected at non-targeted regions in each Panc10.05 surviving colony.**

| sgRNA | Colony 1 | | | | Colony 2 | | | |
|---|---|---|---|---|---|---|---|---|
| | No. of indels | Lowest number of mm | No. of SVs | Lowest number of mm | No. of indels | Lowest number of mm | No. of SVs | Lowest number of mm |
| NT | 424 | 6 | 5 | 9 | 484 | 6 | 13 | 7 |
| NT2 | 483 | 7 | 21 | 8 | 503 | 8 | 16 | 9 |
| 531F(2) | 580 | 7 | 31 | 7 | 420 | 7 | 19 | 8 |
| 52F(3) | 477 | 6 | 26 | 7 | 953 | 6 | 40 | 7 |
| 715F(5) | 534 | 5 | 40 | 9 | 459 | 7 | 31 | 6 |
| 451F(6) | 1156 | 6 | 56 | 7 | 944 | 6 | 48 | 6 |
| 551R(8) | 391 | 7 | 42 | 8 | 453 | 7 | 34 | 6 |
| 164R(14) | 485 | 6 | 72 | 6 | NA | NA | NA | NA |

NA: not available

**Table 5. Breakpoint analyses of surviving colonies from multi-target sgRNA treatment.**

| sgRNA | N | Total bp identified | No. of bp studied* | % of bp studied | indel | 0bp mh (clean break) | 1bp mh | 2bp mh | 3-4bp mh | 5-20bp mh | >20bp mh | Unknown / repetitive |
|---|---|---|---|---|---|---|---|---|---|---|---|---|
| NT(0) | 2 | 25 | 19 | 76% | 2 | 2 | 0 | 3 | 2 | 1 | 2 | 7 |
| 531F(2) | 2 | 58 | 23 | 40% | 5 | 4 | 2 | 2 | 4 | 3 | 2 | 1 |
| 715F(5) | 2 | 42 | 36 | 86% | 4 | 7 | 3 | 6 | 5 | 4 | 7 | 0 |
| 164R(14) | 1 | 31 | 23 | 74% | 8 | 2 | 0 | 3 | 2 | 2 | 3 | 3 |

bp: breakpoint

mh: microhomology

*No. of bp studied: Number of breakpoints in which the junction sequence was studied out of all the breakpoints identified.

**Supplemental Data 1. (separate file)**

List of potential 0-2mm target sites generated by CRISPOR.

Mutation frequency at each site generated by CRISPRessoWGS.

Mutation type determined through IGV.

Data included:

WGS of Panc10.05 surviving colonies mapped to hg19

WGS of selected Panc10.05 surviving colonies mapped to hg38.

(The number of target sites of 176R and 164R differ between hg19 and hg38.)

WGS of TS0111 surviving colonies mapped to hg38
